# Ticks without borders: Microbial communities of immature Neotropical tick species parasitizing migratory landbirds along northern Gulf of Mexico

**DOI:** 10.1101/2023.10.22.563347

**Authors:** Shahid Karim, Theodore J. Zenzal, Lorenza Beati, Raima Sen, Abdulsalam Adegoke, Deepak Kumar, Latoyia P. Downs, Mario Keko, Ashly Nussbaum, Daniel J. Becker, Frank R. Moore

## Abstract

The long-distance, seasonal migrations of birds make them an effective ecological bridge for the movement of ticks. The introduction of exotic tick species to new geographical regions can lead to the emergence of novel tick-borne pathogens or the re-emergence of previously eradicated ones. This study assessed the prevalence of exotic tick species parasitizing resident, short-distance, and long-distance songbirds during spring and autumn at stopover sites in the northern Gulf of Mexico using the mitochondrial 12S rDNA gene. Birds were captured for tick collection from six different sites from late August to early November in both 2018 and 2019. The highest number of ticks were collected in the 2019 season. Most ticks were collected off the Yellow-breasted Chat (*Icteria virens*) and Common Yellowthroat (*Geothlypis trichas*), and 54% of the total ticks collected were from Grand Chenier, LA. A high throughput 16S ribosomal RNA sequencing approach was followed to characterize the microbial communities and identify pathogenic microbes in all tick samples. Tick microbial communities, diversity, and community structure were determined using quantitative insight into microbial ecology (QIIME). The sparse correlations for compositional data (SparCC) approach was then used to construct microbial network maps and infer microbial correlations. A total of 421 individual ticks in the genera *Amblyomma, Haemaphysalis,* and *Ixodes* were recorded from 28 songbird species, of which *Amblyomma* and *Amblyomma longirostre* was the most abundant tick genus and species, respectively. Microbial profiles showed that Proteobacteria was the most abundant phylum. The most abundant bacteria include the pathogenic *Rickettsia* and endosymbiont *Francisella, Candidatus Midichloria,* and *Spiroplasma*. BLAST analysis and phylogenetic reconstruction of the *Rickettsia* sequences revealed the highest similarities to pathogenic spotted and non-spotted fever groups, including R*. buchneri, R. conorii, R. prowazekii, R. bellii, R. australis, R. parkeri, R. monacensis,* and *R. monteiroi*. Permutation multivariate analysis of variance revealed that the relative abundance of *Francisella* and *Rickettsia* drives microbial patterns across the tick genera. We also observed a higher percentage of positive correlations in microbe-microbe interactions among members of the microbial communities. Network analysis suggested a negative correlation between a) *Francisella* and *Rickettsia* and, b) *Francisella* and *Cutibacterium*. Lastly, mapping the distributions of bird species parasitized during spring migrations highlighted geographic hotspots where migratory songbirds could disperse ticks and their pathogens at stopover sites or upon arrival to their breeding grounds, the latter showing means dispersal distances from 421–5003 kilometers. These findings strongly highlight the potential role of migratory birds in the epidemiology of tick-borne pathogens.

Graphic abstract:
Overview of the experimental approach for bird collection and characterization of neotropical ticks microbiome. A) Birds migrating through the Mississippi flyway into the United States were trapped and captured for identification and tick collection and B) immature developmental stages of feeding ticks infesting the birds were carefully removed and stored for morphological and molecular identification of tick species. C) Tick DNA was used for the PCR amplification of arthropods mitochondrial 12S rDNA and amplicons were sequenced and compared to deposited datasets in NCBI database. D) Microbial DNA was isolated using V3-V4 16S primers and unique barcodes attached to amplified 16S sequences prior to Illumina MiSeq sequences. E) Distribution and contribution of birds to tick infestation, and distribution and community profiles of sequenced microbial communities.

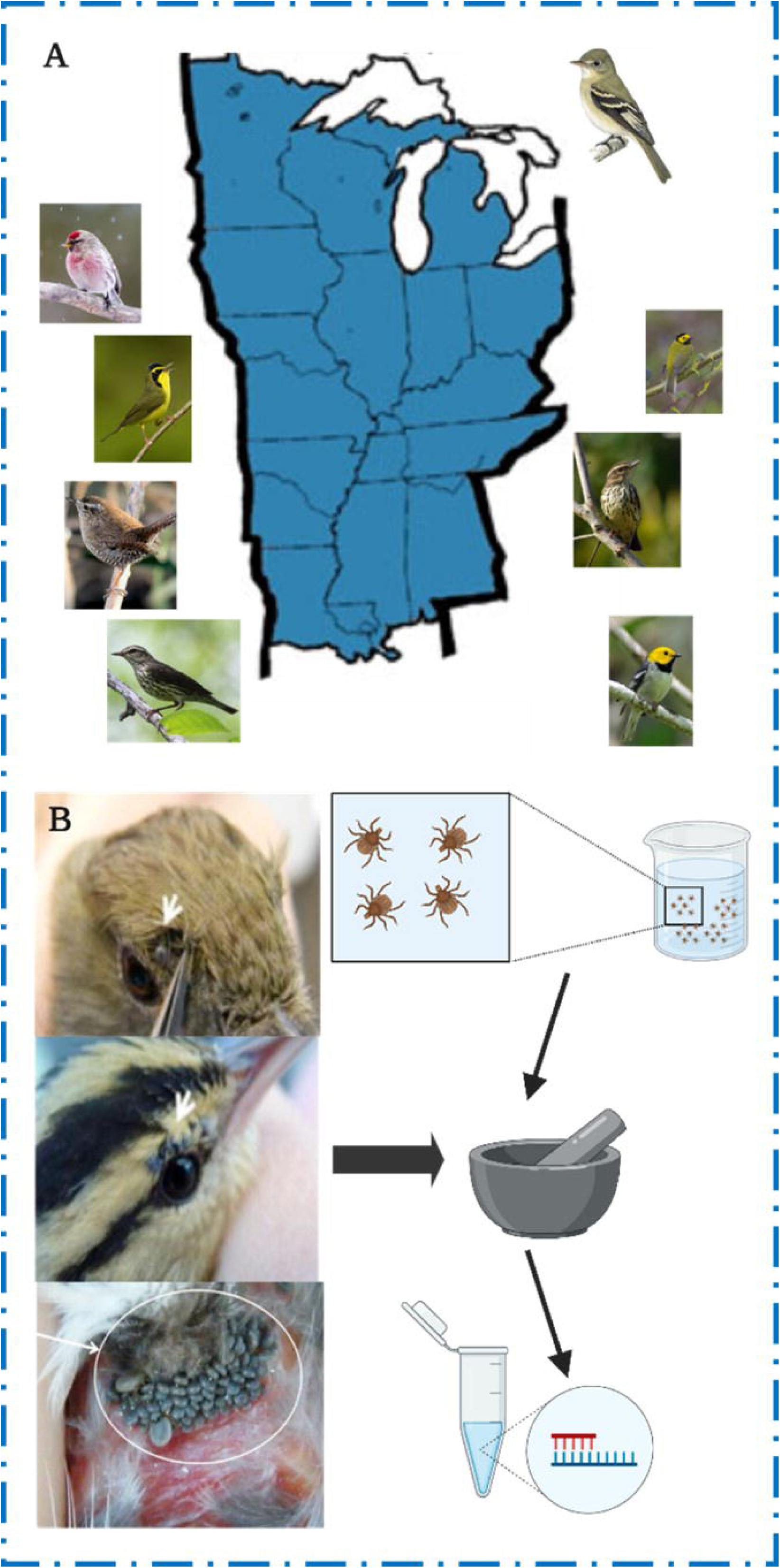

## Introduction

Ixodid (hard-bodied) ticks have a broad host range and are globally distributed, facilitating the transmission of bacterial, viral, and protozoan pathogens that cause diseases in humans and animals (Sonenshine 1993). Transmission of tick-borne pathogens to humans places a significant impact on both public health and the economy, including substantial costs of treatment and disability associated with post-recovery states (Rochlin and Toledo, 2020). In North America, ticks are responsible for over 95% of vector-borne diseases (Rosenberg et al., 2018).

The most prevalent tick-borne illness is Lyme borreliosis, which is caused by the spirochete *Borrelia* (*B.*) *burgdorferi sensu lato* complex and vectored by *Ixodes scapularis*. Over 300,000 and 85,000 cases are reported annually in the USA and Europe, respectively (Lindgren and Jaenson, 2006). Other tick-borne bacterial pathogens found in the USA include but are not limited to *Rickettsia rickettsii, Anaplasma phagocytophilum, R. parkeri, Ehrlichia chaffeensis*, and *E. ewingii*. Tick-borne viral pathogens belong to the families *Flaviviridae, Bunyavirales, Orthomyxoviridae*, and *Reoviridae*. One emerging tick-borne viral pathogen is Powassan virus (POWV), which was first reported in 1958 in Powassan, Ontario (McLean et al., 1962). Tick-borne flaviviruses are geographically prevalent in Europe, Russia, China, and Japan (Mansfield et al., 2017; Rochlin and Toledo, 2020). However, POWV is endemic in the northeast and upper midwest of the USA, though cases have also been reported in far-eastern Russia (Anderson and Armstrong, 2012). The emergence of the invasive Asian long-horned tick, *Haemaphysalis longicornis*, in the USA (Rainey et al., 2018; Rochlin, 2019) poses a potential threat to the livestock industry, as it is the competent vector of *Theileria orientalis*, the protozoan that causes theileriosis (Fujisaki et al., 1994; Hammer et al., 2015)., which is characterized by anemia and occasional mortality.

Although terrestrial mammals commonly serve as hosts for ticks, they typically travel relatively short distances before ticks complete their blood meal and drop off (Cheng 1967). However, ticks of the genera *Argas, Ornithodoros, Ixodes, Amblyomma,* and *Haemaphysalis* also can parasitize birds (James et al., 2011; Barros-Battesti et al., 2006; Hasle et al., 2011; Movila et al., 2011; Zeringota et al., 2017). In the case of migratory avian hosts, ticks can disperse much further from their origin. Many species of migratory songbirds, for example, fly long distances, sometimes moving across continents and extensive geographical features (e.g., oceans, deserts) within a relatively short period every spring and autumn (Deppe et al., 2015; Ouwehand and Both 2016). Due to the energetic cost of migration (Wikelski et al., 2003; McWilliams et al., 2004), most migratory birds must stop *en route* at stopover sites to rest and replenish fuel stores in unfamiliar habitats. Birds may pick up new or drop off existing ectoparasites, such as ticks, during stopover, acting as a long-distance dispersal mechanism for ectoparasites and their pathogens (Mukherjee et al., 2014; Cohen et al., 2015; Zenzal et al., 2017; James et al., 2011; Hasle 2013; Nicholls and Callister 1996; Becker et al., 2022). Moreover, migratory birds often serve as competent reservoirs of multiple tick-borne pathogens (Reed et al., 2003; Comstedt et al., 2006; Becker and Han, 2021) and thus are rightfully termed as "ecological bridges" due to their role in moving both ectoparasites and pathogens to new ecological niches (Pascucci et al., 2019).

Exotic ticks transported into North America by migratory birds have thus far had limited success in becoming locally established, which may stem from the unavailability of suitable climatic conditions and/or natural host species as well as competition with native ticks (Hasle 2013). If ticks transported by migratory birds were to succeed in establishing permanent populations in North America, the pathogens they carry may start infecting local native hosts and tick species. With a changing climate, non-native tick species dispersed by migratory birds could eventually establish populations in the USA with the potential of becoming invasive.

Invasive species cost over $26 billion per year in the USA in economic, environmental, and public health damages (Crystal-Ornelas et al., 2021). One example is the recent invasion of *Haemaphysalis (H) longicornis,* the Asian long-horned tick. This invasive species was first identified in New Jersey in 2017 (Rainey et al., 2018) and was then confirmed in at least 14 other states (USDA APHIS 2021; Tufts et al., 2019). *Haemaphysalis (H) longicornis* is native to East Asia, and geographic expansion could present a significant threat to animal and public health by serving as a competent vector of pathogens including *Theileria, Babesia, Rickettsia*, and severe fever with thrombocytopenia syndrome virus (Thompson et al., 2020; Luo 2015; Fill et al., 2017; Godsey et al., 2016). Additionally, *H. longicornis* is able to reproduce parthenogenetically (without a male), furthering the potential spread and impact of the speices.

This study aimed to investigate the diversity of commensal and pathogenic bacteria within the microbial community of Neotropical ticks collected from songbirds during spring and autumn when many of them stopovers along the northern Gulf of Mexico coast when entering or leaving the USA. We wanted to determine if ticks infesting birds can serve as sentinels to monitor the introduction of known and previously unreported pathogenic bacteria into new geographical areas. In this study, primarily migratory birds were captured at stopover sites along the northern Gulf of Mexico coast during spring and autumn migration. Ticks were collected from these migratory birds, and a fragment of each tick’s mitochondrial 12S rDNA gene sequence was amplified to identify species. A high throughput 16S rRNA sequencing approach was used to examine the microbiome of individual immature ticks to determine their microbial community structure, composition, and stability. Lastly, we assessed geographic overlap in the migration and breeding distributions of songbirds found to be carrying ticks during spring migration to identify spatial hotspots of potential tick and pathogen dispersal capacity into the USA.

## Materials and Methods

### Ethics approval

All animal sampling was conducted in strict accordance with the recommendations in the Guide for Care and Use of Laboratory Animals of the National Institutes of Health, USA. Collection and handling of birds were approved by the U.S. Geological Survey Bird Banding Laboratory (permit #24101), the Louisiana Department of Wildlife and Fisheries, the Alabama Department of Conservation and Natural Resources, and the Institutional Animal Care and Use Committee (IACUC) at the University of Southern Mississippi (protocol #17081101) and the U.S. Geological Survey (protocol #LFT 2019- 05).

### Materials

All common laboratory supplies and chemicals were purchased through Bio-Rad (Hercules, CA, USA), Sigma-Aldrich (St. Louis, MO, USA), and Fisher Scientific (Grand Island, NY, USA), unless specifically noted.

### Bird capture and sampling

Songbirds were sampled at six stopover sites along the northern Gulf of Mexico during autumn (late August to early November) in 2018 and 2019 and seven stopover sites during spring (mid-March to mid-May) in 2018 and 2019. Birds were captured using 15– 20 nylon mist nets (12 or 6 x 2.6 m; 30-mm mesh) per site from up to four sites in southwest Louisiana and three sites in southern Alabama (Table 1). Netting was conducted daily with the exception of inclement weather. During spring, nets were operated from local sunrise until approximately five hours after sunrise; during three days of the week, nets were reopened from three hours before local sunset until ∼30 minutes before sunset. During autumn, nets were operated from local sunrise until approximately six hours after sunrise. Each captured individual was ringed with a uniquely numbered U.S. Geological Survey metal leg band, had its condition assessed, morphometrics recorded, and, when time allowed, inspected for the presence of ectoparasites. Bird species were categorized as resident, short-distance migrants, or long-distance migrants based on where the species predominantly spends the stationary non-breeding period. Resident species were found year-round at our study sites, short-distance migratory species spent the stationary non-breeding season north of the Tropic of Cancer, and long-distance migratory species spent the stationary non-breeding season south of the Tropic of Cancer (DeGraaf and Rappole 1995; Carlisle et al., 2004; Zenzal et al., 2018). When a tick was discovered attached to a bird, we collected the tick in the field by carefully detaching it with fine-tipped forceps and preserved it in a vial of 70% ethanol. For each tick sample, we recorded the collection date, unique sample code, and the bird’s unique leg band number.

**Table 1:** Bird trapping and tick collection sites including state and coordinates.

| *Site* | *State* | *Coordinates* |
| --- | --- | --- |
| *Grand Chenier* | Louisiana (LA) | 29.749659, -92.894811 |
| *Grand Chenier* | Louisiana (LA) | 29.738521, -92.836370 |
| *Hayes* | Louisiana (LA) | 0.109666, -92.908688 |
| *Reeves* | Louisiana (LA) | 30.435878, -93.075420 |
| *Gulf Shores* | Alabama (AL) | 30.260475, -87.748360 |
| *Saraland* | Alabama (AL) | 30.811519, -88.058210 |
| *Perdido* | Alabama (AL) | 31.024289, -87.677632 |

### Tick collection and molecular identification

Ticks collected from songbirds were mostly immature developmental stages. Taxonomic keys for immature stages of exotic or uncommon ticks are not available for all species. Furthermore, engorged immature specimens are often damaged to such an extent that diagnostic morphological characters are obliterated. Therefore, individual whole ticks were molecularly identified by using 12S DNA sequencing. A fragment of each tick’s mitochondrial 12S rDNA gene sequence was amplified from 485 samples with T1B and T2A primers (Beati & Keirans, 2001) obtained from Invitrogen. Dream Taq Polymerase (Invitrogen, Thermo Fisher Scientific, Waltham, MA, USA) was used to amplify tick DNA. The amplicons were bidirectionally sequenced at Eurofins Genomics (Eurofins Genomics, Louisville, KY, USA). Complementary strands were visually examined and assembled by using Benchling. The sequences were compared to homologous nucleotide fragments in GenBank using BLAST (Mukherjee et al., 2014).

### Statistical analysis of tick infestation

To analyze tick infestation data, we used generalized linear mixed models (GLMMs) via the *lme4* package in the R statistical language (Bates et al., 2015; R Core Team 2023). We used a two-step process to quantify species-level variation in tick outcomes and to assess additional temporal and migratory drivers of infestation. For binary tick positivity across all birds sampled during our study (*n* = 17,550), we first fit a GLMM with a fixed effect of bird species and a site-level random effect to account for regional artifacts. We then had another GLMM with an additional random effect for bird species that included fixed effects of year, migratory season, and their interaction. To analyze the intensity of tick parasitism on birds with identified ticks (*n* = 164), we likewise fit a GLMM with Poisson errors and an observation-level random effect nested within the site random effect to account for overdispersion (Harrison 2014). We then fit another equivalent GLMM with an additional random effect for bird species that included fixed effects of year, migratory season, and migratory category. We excluded bird species with only one tick association (*n* = 7 species) from these intensity GLMMs. We assessed Poisson GLMMs for overdispersion and adjusted any post-hoc comparisons from our models using the Benjamini–Hochberg correction. We derived model fit with marginal and condition R^2^ (Nakagawa and Schielzeth 2013).

### Library preparation for Illumina 16S sequencing

Sixty-two of the 421 genomic DNA isolated from individual ticks did not pass quality control (QC) and the 359 that passed QC were used to generate libraries according to the library generation protocol by Illumina Indexing Methodology (Kumar et al., 2021). Briefly, a two-stage PCR amplification process was used to amplify the 16S rRNA V3- V4 region, followed by a dual indexing step that assigns unique index sequences to the V3-V4 amplicons. The concentration of the resulting PCR products were determined using qPCR and equal concentration of each sample was pooled and sequenced in a single run of an Illumina MiSeq sequencing instrument using reagent kit v2 (500 cycles) with 2 × 250 bp output at the University of Mississippi Medical Centre (UMMC) Genomics Core Facility. DNA extraction controls, PCR controls, and known mock bacterial communities (ZymoBIOMICS™ Microbial Community DNA Standard, Irvine, CA, USA) were simultaneously processed alongside the tick samples. All critical steps including determination of amplicon size, and amplicon purification following each PCR step were performed as described by Kumar et al. (2021).

### Tick microbiome data processing

Unless otherwise stated, all data preprocessing was done following the video tutorial of the Quantitative Insights into Microbial Ecology (QIIME2) pipeline (Bolyen et al., 2019). Briefly, demultiplexed fastq files were unzipped and the forward and reverse fastq files merged into a single fastq file using Casava (insert version here). The Atacama soil microbiome pipeline was incorporated to control demultiplexed paired-end reads using the DADA2 plugin as previously described (Callahan et al., 2016). Low-quality and Chimeric sequences were removed; subsequent merging of paired-end reads ensured 20 nucleotide overhangs between forward and reverse reads. We accomplished taxonomic assignment and diversity metrics in QIIME2 as described (Quast et al., 2013). Raw sequences will be submitted to the required databases.

### Tick microbiome visualization

Microbiome Analyst, a web-based interface, was used for data visualization by employing taxonomy and metadata tables generated from data processing as input files (Dhariwal et al., 2017; Chong et al., 2020). Low count and prevalence data were filtered from the OTU table by setting values of 10 and 20, respectively. A filtered abundance table was exported and used in generating histograms of bacterial abundance in Microsoft Excel 2016 (Microsoft, 2018). Network correlation maps were constructed based on the sparse correlations for compositional data (SparCC) approach (Friedman and Alm, 2012). This approach uses the log-transformed values to carry out multiple iterations, which subsequently identifies taxa outliers to the correlation parameters (Chong et al., 2020). To compare the differences in the microbiome between tick groups based on measures of distance or dissimilarity, a matrix was generated from log-transformed sequence data, using the Bray-Curtis distance matrix and ordination of the plots was visualized using Principal Coordinates Analysis (PCoA).

### Statistical analysis of tick microbial communities

Kruskal–Wallis tests followed by Dunn’s multiple comparison tests were used to compare the differences in alpha diversity between all identified ticks at the genus and species level based on the observed OTU metric and Shannon’s diversity index. Permutational multivariate analysis of variance (PERMANOVA) was used to determine significant pairwise differences in the tick microbial communities by comparing the means across different tick genus and species. Statistically significant data were represented as P <0.05.

### Spatial analysis of parasitized birds

To identify spatial hotspots of potential tick and pathogen dispersal into North America, we mapped the distributions of bird species parasitized by ticks during spring migration, when detected ticks most likely originated from Central and South American wintering grounds. For those parasitized migratory birds, we aggregated species shapefiles from the International Union for Conservation of Nature using the *rgdal* and *rgeos* packages in the R statistical language (Zedan, 2004). Aggregated avian distributions were stratified by migration and breeding seasons to illustrate dispersal capacity to stopover sites and the breeding grounds respectively, for each identified tick species (Zedan, 2004). Lastly, we calculated distance between spring capture site and the centroid of the breeding range for each parasitized bird species using the *geosphere* package, for each unique combination of site, bird species, and tick species. We then used a GLM with a Gamma distribution to identify how mean dispersal capacity during spring migration varied across bird-infesting tick species.

## Results

### Tick infestation and identification

A total of 164 individual birds of 28 species were found with attached ticks during our spring and autumn sampling periods (Table S1). When considering tick parasitism status across the 17,550 birds sampled, bird species did not significantly differ in their odds of infestation (χ^2^ = 104.85, df = 103, P = 0.43, R^2^_m_ = R^2^_c_ = 0.22), likely owing to the overall low prevalence of parasitism (<1%). After accounting for site and species variation, tick positivity varied significantly by sampling year and migratory season (interaction: χ^2^ = 5.96, P = 0.01, R^2^_m_ = 0.19, R^2^_c_ = 0.20). The odds of birds harboring ticks were greatest in spring 2019 compared to all other sampling periods (*z* > 5.2, P < 0.01).

Tick intensities among parasitized birds were highly variable across host species (Fig. 1; χ^2^ = 33.93, df = 20, P = 0.03), which explained 17% of the variance (R^2^_m_) compared to the individual-and site-level random effects (R^2^_c_ = 0.54). The Yellow-breasted Chat (*Icteria virens*) and Common Yellowthroat (*Geothlypis trichas*) had the greatest tick intensities (x□ = 18 and 11), although only one individual per species was parasitized. Most ticks were contributed by the Hooded Warber (*Setophaga citrina*, 31%) and Swamp Sparrow (*Melospiza georgiana*, 16%), whereas the least ticks were contributed by the Red-eyed Vireo (*Vireo olivaceus*, <1%), Scarlet Tanager (*Piranga olivacea*, <1%), White-throated Sparrow (*Zonotrichia albicollis*, <1%), Winter Wren (*Troglodytes hiemalis*,<1%), American Redstart (*Setophaga ruticilla*, <1%), and Northern Waterthrush (*Parkesia noveboracensis*, <1%). Most ticks (54%) were collected at the stopover site in Grand Chenier, LA.

**Figure 1:**
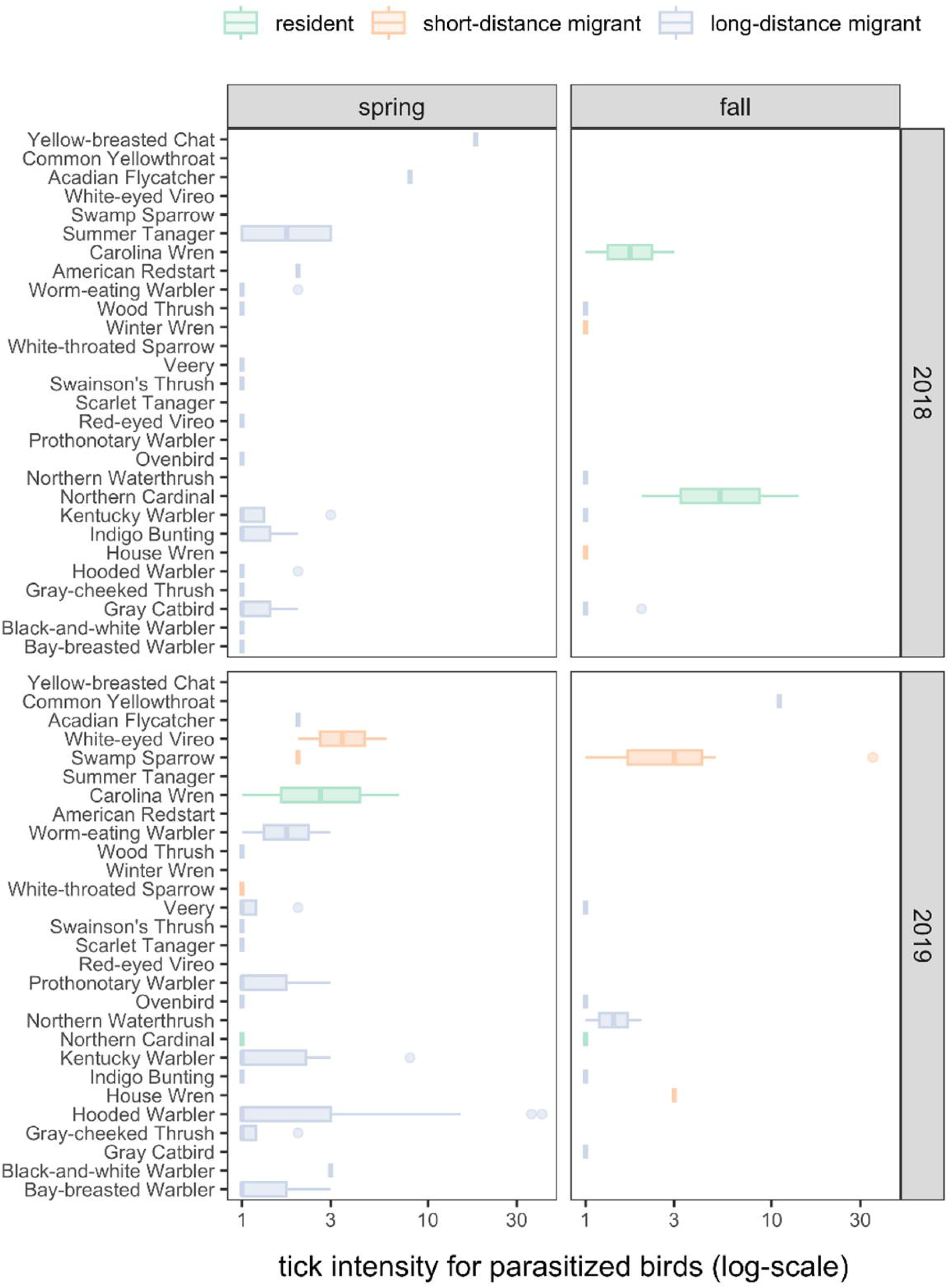
The distribution of tick intensities across bird species, years, and seasons

Our secondary Poisson GLMM (R^2^_m_ = 0.08, R^2^_c_ = 0.52) revealed no significant variation in tick intensity by migratory season (χ^2^ = 0.28, P = 0.60) or year (**χ^2^** = 2.38, P = 0.12), although birds tended to have more ticks in 2019 (x□ = 3.09) and during autumn migration (x□ = 3.00). Tick intensities did vary by migration category (χ^2^ = 7.24, P = 0.03), which was driven by short-distance migrants having greater tick intensities on average than long-distance migrants (x□ = 4.15 vs x□ = 2.24; z = 2.36, P = 0.05), neither of which differed significantly from residents (x□ = 3.44; short-distance migrants: z = 0.22, P = 0.82; long-distance migrants: z = 1.82, P = 0.10). Because short-and long-distance migratory birds were more commonly sampled, these species contributed 73% and 20% of all identified ticks (Fig. 1). Most ticks were collected in 2019 (71%) and during spring migration (70%). Neither of our tick intensity GLMMs showed significant overdispersion (χ^2^ 18.74, P = 1).

Of the 421 tick samples collected from songbirds, 62 did not yield DNA of sufficient quality for amplification and sequencing. The remaining 359 specimens were identified as *Amblyomma americanum, A. maculatum, A. calcaratum, A. coelebs, A. geayi, A. longirostre, A. nodosum, A. ovale, A. parvum, A. sabanerae, A. triste, A. varium, Haemaphysalis leporispalustris, Ixodes brunneus, I. dentatus,* and *I. scapularis*. Four tick species accounted for 81% of all ticks collected during the study period, with *A. nodosum* (29%; 120/421,) *A. longirostre* (20%; 83/421)*, H. leporispalustris* (16%; 66/421), and *A. maculatum* (16%; 69/421) representing the most tick species, respectively (Table S1).

### Tick DNA sequencing results

Illumina 16S rRNA sequencing of all 359 tick DNA samples produced 11,064,738 raw forward and reverse reads. The distribution of the raw reads ranges from an average of 26,957 reads per sample and a maximum of 89,223 reads per sample (Table S2). Rarefaction analysis of the individual samples from a sequencing depth of 500 to 5,000 confirmed there was adequate sequence coverage relative to the number of observed features and a plateau for individual tick samples (Fig. S1).

### Bacterial abundance and distribution

Overall, 1,416 unique bacterial OTUs were identified across each tick genera (Table S3). Proteobacteria represented the most abundant phylum across the three genera of ticks sequenced. The abundance of the phylum proteobacteria ranges from 82% to 98% of the total bacterial OTU (Fig. S2A). Other identified OTUs were either classified in the phyla Actinobacteriota, Firmicutes, Cyanobacteria, or unassigned to any known bacteria phyla. At the genus level, irrespective of tick species, *Francisella* was the most abundant bacteria (60-75%), and represents the only bacteria genus identified in *A. varium* (Fig. 2; Fig. S2B). Including *Francisella*, 10 of 11 bacteria have reads greater than 1% in at least one tick (Fig. 2; Fig. S2B). *Rickettsia* had the second highest relative abundance (11-14%) in all ticks, while *Spiroplasma* and *Cutibacterium* were present in relatively high abundance in *Haemaphysalis* and *Ixodes* ticks respectively (Fig.2; Fig. S2B). *Coxiella* was present in *A. coelebs* (5%), *A. calcaratum* (13%), and *A. geayi* (60%), while *Candidatus Midichloria* was present in *A. maculatum* (5%) *I. brunneus* (37%).

**Figure 2:**
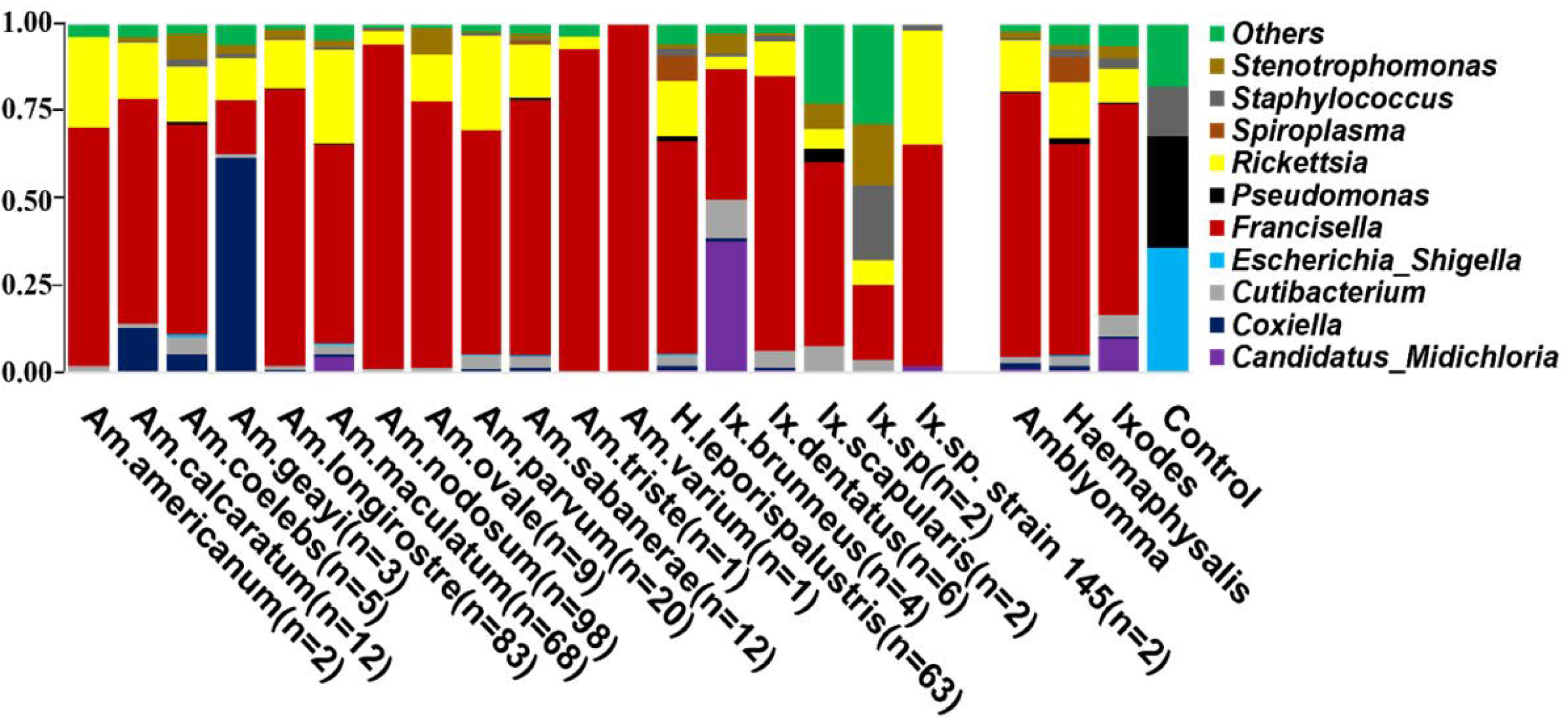
Relative abundance of the top 10 bacterial genera identified from ticks collected off migratory birds. Each horizontal bar represents the cumulative average of the bacterial abundance identified from each tick species and genus.

### Tick bacterial diversity

Ticks from the genus *Amblyomma* exhibited the lowest bacterial alpha diversity, while *Haemaphysalis* and *Ixodes* have similar levels of alpha diversity (Observed OTUs: χ^2^ = 36.34, P < 0.001; Shannon: χ^2^ = 66.75, P < 0.001; Fig. 3A). *A. coelebs* had the highest bacterial alpha diversity, while *A. nodosum* had the least diverse bacterial communities based on observed OTUs (χ^2^ = 59.8, P < 0.001) and Shannon’s diversity index (χ^2^ = 65.776, P < 0.001) when compared to all other *Amblyomma* species (Fig. 3B). No difference in alpha diversity was observed within *Ixodes* and *Haemaphysalis* species (Fig. S3).

**Figure 3:**
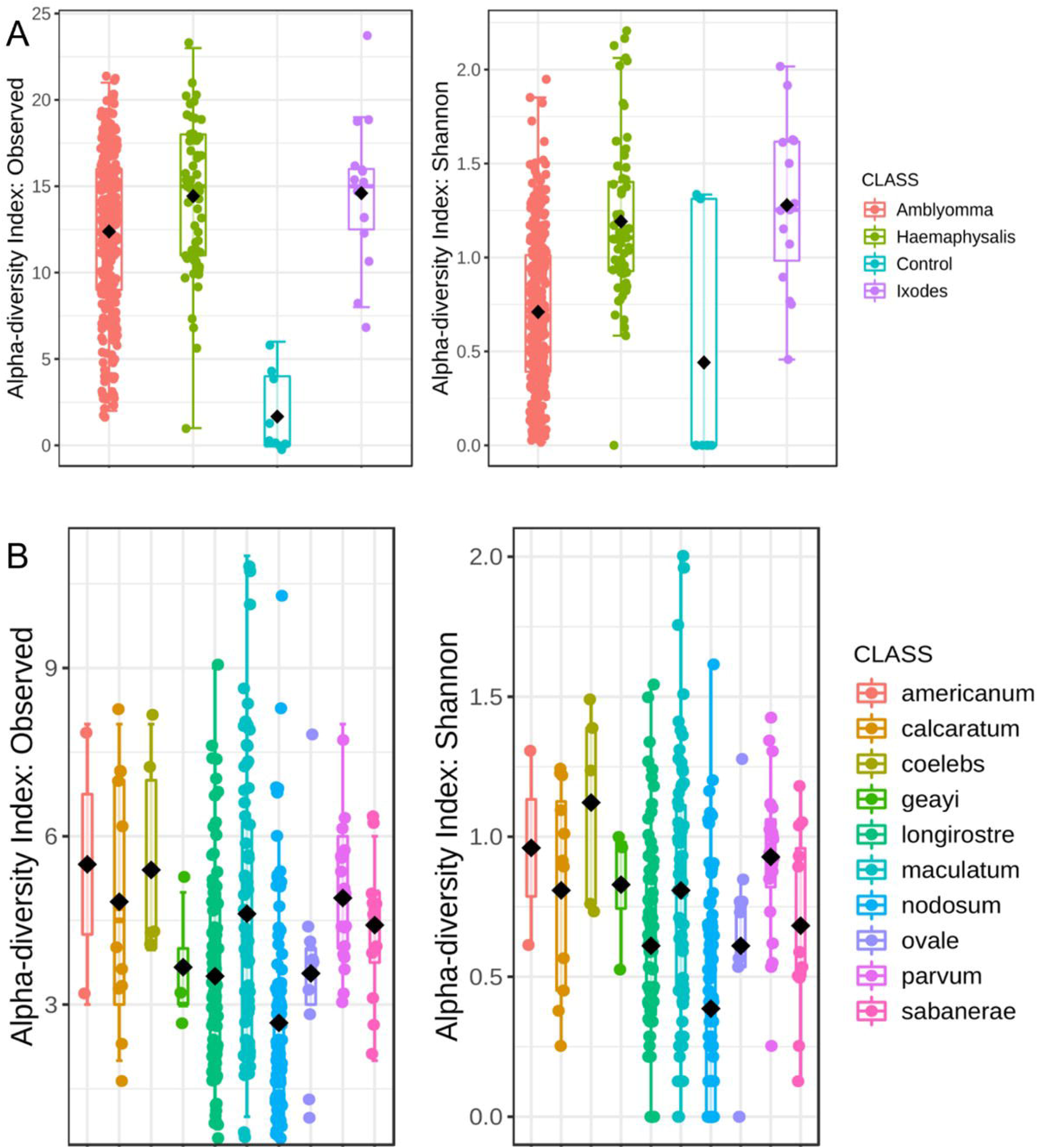
Measures of alpha diversity for A) combined tick genera and B) *Amblyomma* species. Alpha diversity was estimated using the observed OTUs and Shannon’s Index following rarefaction analysis of samples to a depth of 5000 sequence.

Our analysis of microbial communities revealed significant differences in the bacterial communities between all tick genera (*Amblyomma, Ixodes,* and *Haemaphysalis*). The NMDS analysis of community structure using the Bray-Curtis distance matrix (PERMANOVA: F = 30.04, R^2^ = 0.23, P < 0.001, [NMDS] Stress = 0.11) revealed overlapping clusters across all tick genera (Fig. 4A; Fig. S4A). Several *Amblyomma* and *Haemaphysalis* ticks were seen to cluster separately on the lower axis of the PCoA (Fig. 4A).

**Figure 4:**
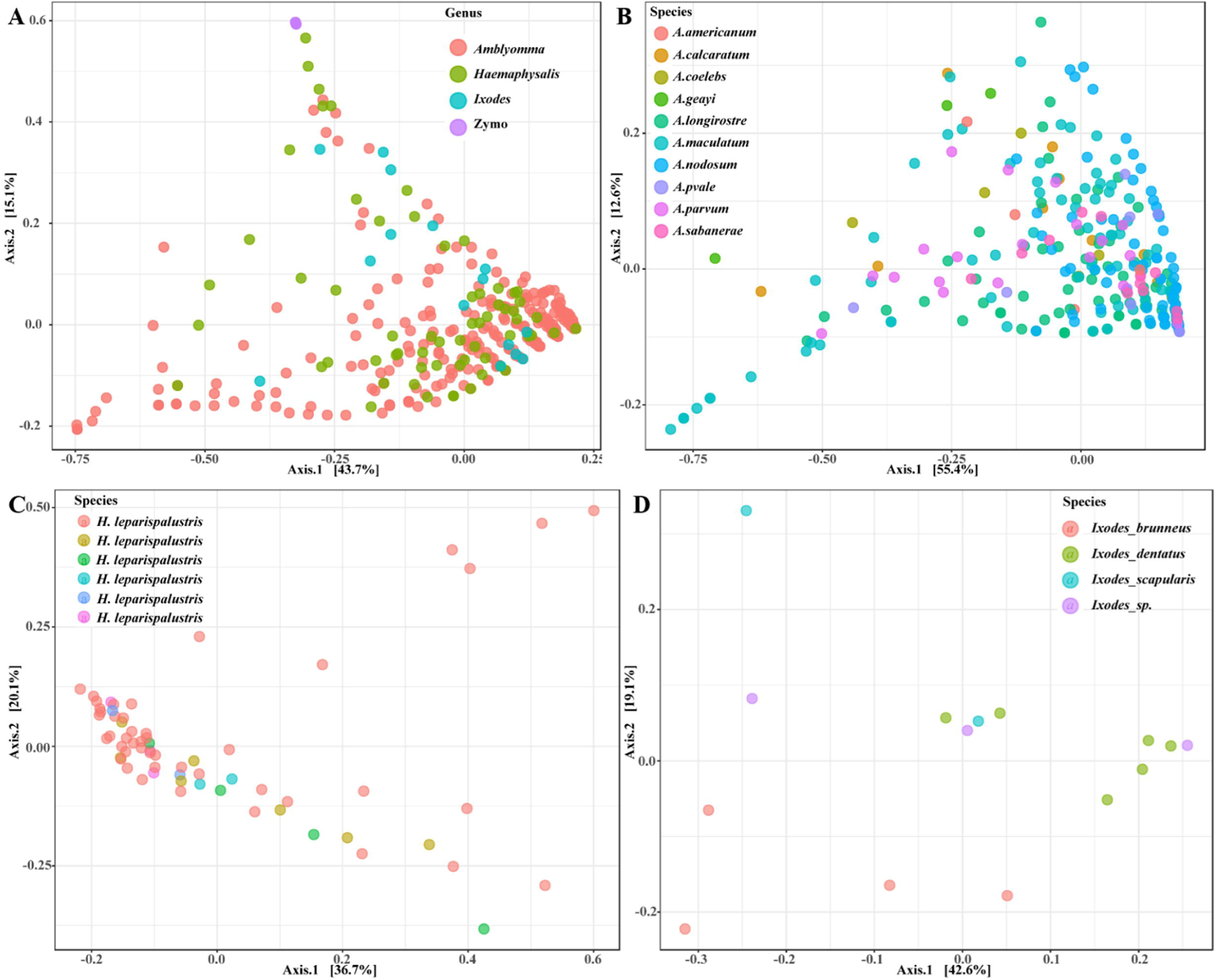
Principal coordinate analysis of beta diversity measures for A) all tick genus,B) *Amblyomma* C) *Ixodes* and D) *Haemaphysalis leporispalustris* ticks using the Bray-Curtis similarity metrics. Highlighted ellipses indicate measures of confidence based on degree of variations.

**Figure 5:**
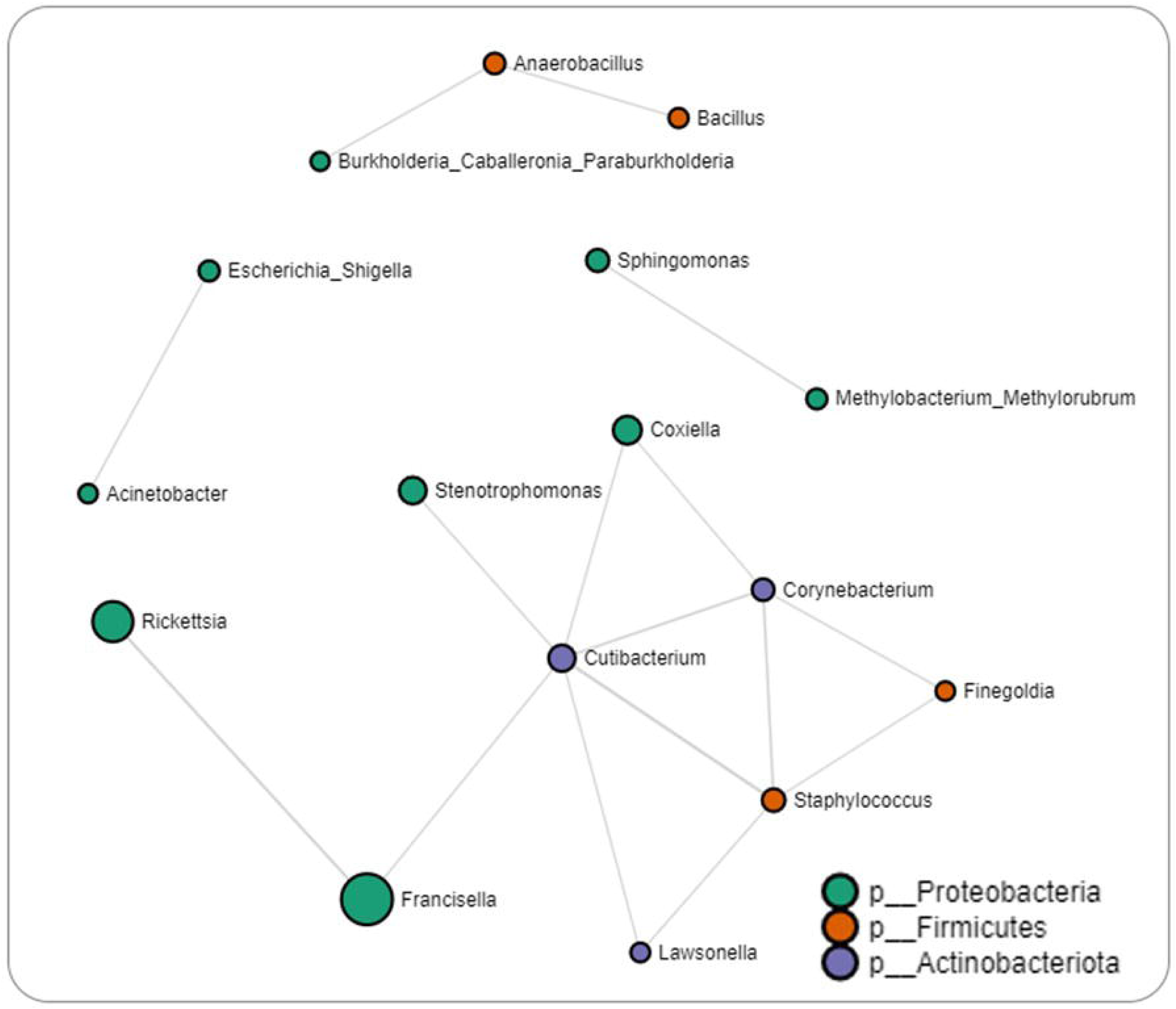
Correlation network analysis on all tick genera. Correlation network generated using the SparCC algorithm. Correlation network with nodes representing taxa at the family level and edges representing correlations between taxa pairs. Node size correlates with the number of interactions in which a taxon is involved. The color-coded legend shows the bacterial phyla.

We observed differences in the abundance of *Francisella* and *Rickettsia* between ticks clustered together and those widely dispersed (Fig. S4B and S4C). While most of the *Amblyomma* species tend to share similar microbial communities based on the PERMANOVA (F = 10.33, R^2^ = 0.24, P < 0.001), several *Amblyomma* ticks, in particular *A. maculatum, A. calcaratum, A. geayi,* and *A. longirostre,* were outliers (Fig. 4B). The differences in the respective abundance of *Rickettsia* and *Francisella* were responsible for the clustering patterns observed in the *Amblyomma* ticks and for the *Amblyomma* species clustered as outliers (Fig. S5A and S5B). *Ixodes* ticks differed significantly according to the PERMANOVA of the Bray-Curtis distances (F = 2.60, R^2^: 0.41, P < 0.004). Distinct clustering was observed with *I. dentatus, I. scapularis* and *I. sp*, while *I. brunneus* clustered separately as explained by 18.2% (Axis 1) and 44.8% (Axis 2) of the PCoA (Fig. 4C; Fig. S5C and S5D). Unlike the *Amblyomma* and *Ixodes* ticks, only one *Haemaphysalis* species was part of this study, and no unique clustering was observed in this specie. Similarly, the PERMANOVA test of the Bray-Curtis distances was not statistically significant (Fig. 4D).

### Tick microbial interactions

Our network analysis identified 16 significant partial correlations between 12 OTUs. 87% of interactions were identified as positive correlations (Table S4). Negative partial correlations were observed between the abundance of the genera *Francisella* and *Rickettsia* as well as the genera *Francisella* and *Cutibacterium*, which were the only two bacteria with which *Francisella* interacted. Within the network interactions, four unique clusters were identified with the largest clusters consisting of bacteria commonly reported within the tick microbiome such as *Coxiella, Francisella, Rickettsia, Staphylococcus* and *Cutibacterium* (Fig. 7). Log-transformed counts of *Rickettsia, Francisella,* and *Coxiella* showed significantly higher abundance in *Amblyomma* ticks (Fig. S6). At the genus level, 65 microbe–microbe interactions were observed in the *Amblyomma* network, 54 in the *Ixodes* network, and 48 in the *Haemaphysalis* network. Detailed microbe-microbe interactions observed amongst each tick genus can be found in the additional file section (Table S5-S9).

### Spatial analysis of parasitized birds

A total of 17,550 birds (comprising 14,929 unique individuals) were captured and sampled for ticks, encompassing 101 bird species. Among them, 164 individual birds, representing 28 bird species, were found to have ticks. Of the 28 bird species, we focused our spatial analyses on the 21 species of migratory birds sampled in spring for which ticks could be molecularly identified to species. Geographies of plausible tick and pathogen dispersal varied widely among the 12 tick species (Fig 6). This analysis suggested migratory birds using the Gulf of Mexico during spring migration could disperse *Amblyomma calcaratum*, *A. coelebs*, *A. longirostre*, *A. nodosum*, *A. parvum, and A. varium* possibly as far northwest as Alaska, whereas *A. geayi*, *A. ovale*, *A. triste*, and *Haemaphysalis leporispalustris* had less dispersal capacity (e.g., north into the Great Lakes region) owing to the shorter-distance migrations of their avian hosts.

**Figure 6:**
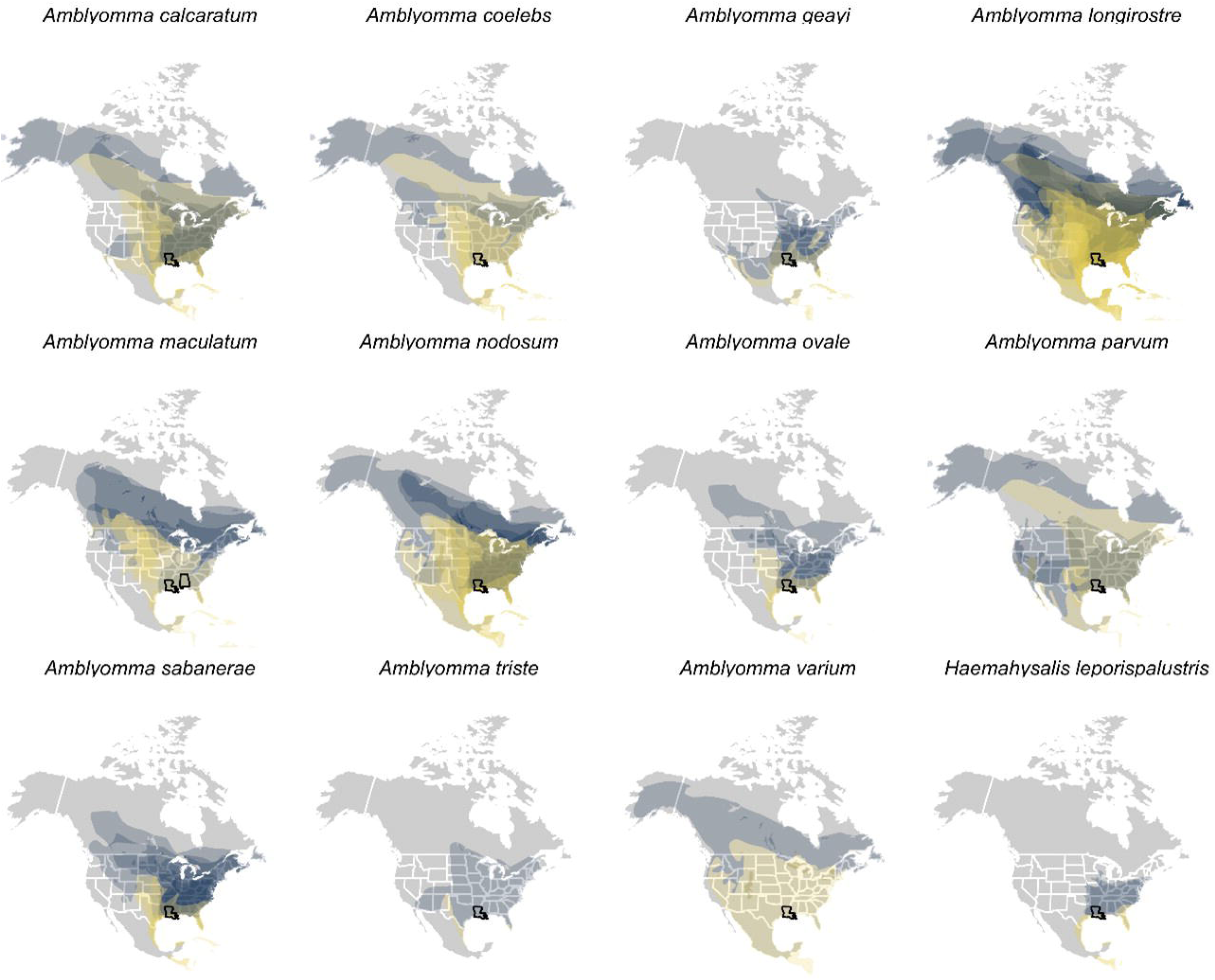
International Union for Conservation of Nature (IUCN) distributions of those bird species and prediction of their potential spread. Each plot is for a different tick species, so some ticks are obviously found on more species than others. The IUCN distributions includes the breeding range (blue) and the migration range (yellow), a prediction of where the migratory birds could spread ticks during stopover (yellow) or upon their arrival to the breeding area (blue).

Formal analysis of distances between spring migration capture sites and breeding range centroids for parasitized birds (n = 69 unique combinations of bird species, tick species, and sites) demonstrated dispersal capacity ranged from 421 to 5003 kilometers (Fig 7). On average, *A. geayi* had the least dispersal capacity (x□ = 925), whereas *A. coelebs* had the most extreme dispersal capacity (x□ = 3,331 km). A GLM excluding the two tick species with only one parasitized bird species or capture site (*A. triste* and *A. varium*) suggested such dispersal capacity among tick species to be approaching significance (χ^2^ = 15.52, P = 0.08).

**Figure 7:**
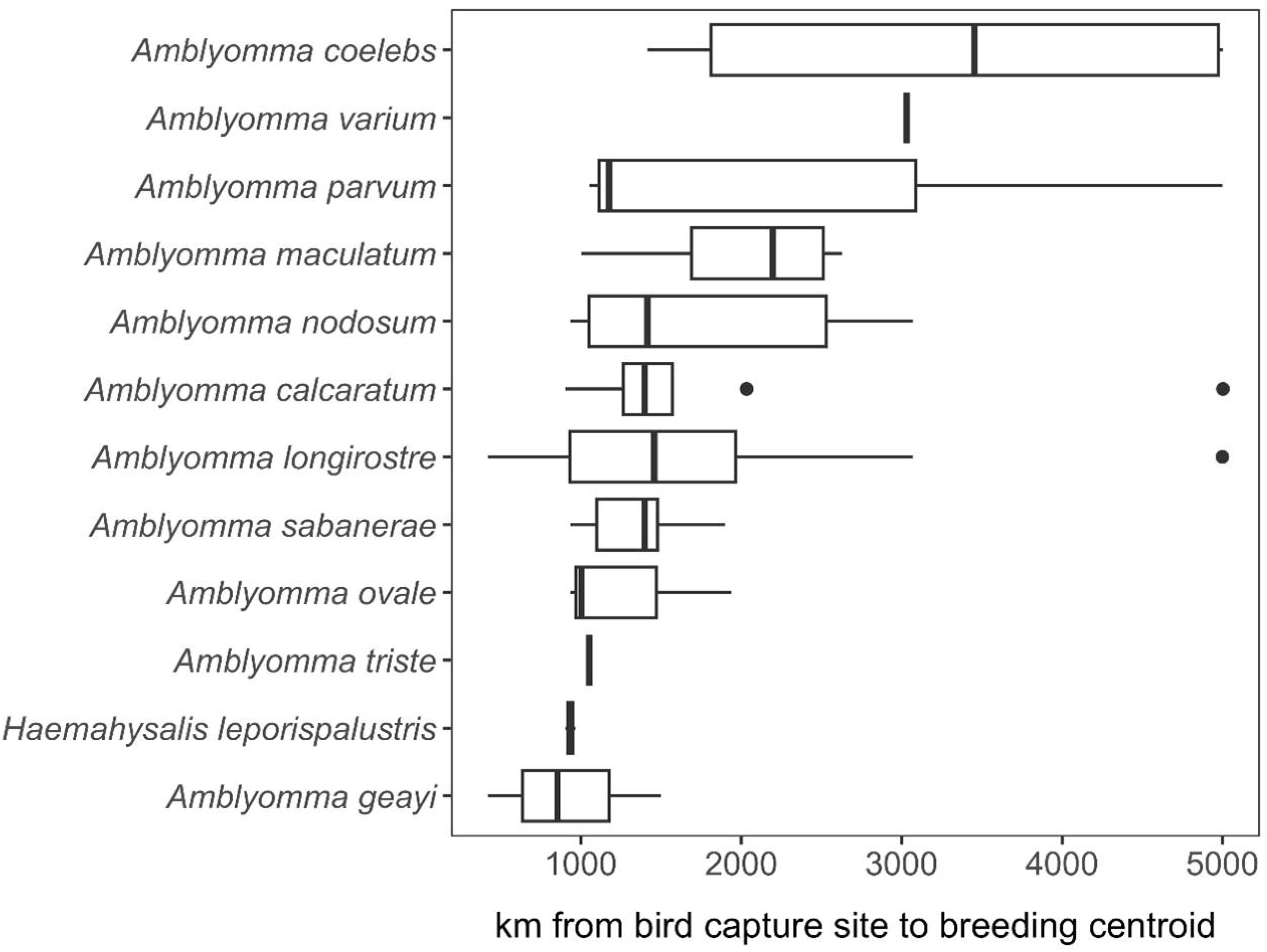
Estimated dispersal distance of each tick species recovered from migratory birds in spring.

## Discussion

This study aimed to assess the role of birds in introducing exotic neotropical tick species and their associated microbes from Central and South America into North America. The results of this study indicate that bird-associated factors, such as the distance with which they migrate, can play important roles in introducing exotic tick species into new areas. Metagenomic sequencing also revealed significant diversity in the microbial communities within the ticks, members of which include bacteria in the pathogenic and symbiotic genera.

### Prevalence of bird infestation and seasonality

We found the most ticks in late spring (April), when over 65% of the ticks were collected, although ticks were collected over the course of spring and autumn.

### Microbial diversity in exotic tick species

In the current study, we aimed to characterize the microbiome of ticks collected from birds migrating to and from the USA, since they can disperse tick species and their associated pathogens. Our previous work during spring migration in Louisiana identified Neotropical tick species including *Haemaphysalis juxtakochi*, *Amblyomma longirostre*, *A. nodosum*, *A. calcaratum*, *A. maculatum*, and *H. leporispalustris* (Budachetri et al., 2017). The identified ticks were exotic species that originated outside of the USA (Budachetri et al., 2017). Our 12S rDNA analysis here identified three tick genera— *Amblyomma, Ixodes,* and *Haemaphysalis*—based on the homology search. These findings further support the role of migratory birds in the introduction and dispersal of ticks in the general *Haemaphysalis* and *Amblyomma* as previously reported (Budachetri et al., 2017). The endosymbiont *Francisella* accounted for more than half of the bacterial abundance detected from each tick species, followed by *Rickettsia*. *Francisella* was the only identified genus in *A. varium*. The presence of *Coxiella* in three *Amblyomma* species and *Candidatus Midichloria* in *Ixodes brunneus* was correlated with lower abundance of *Francisella*. The high association of *Francisella* with all tick species tested suggests its presence as an obligate endosymbiont, although the presence of *Rickettsia* and *Candidatus Midichloria* would also suggest that these genera play a crucial role in the development of these tick species. Both *Candidatus Midichloria* and *Francisella*-like endosymbionts have been most commonly reported in ixodid ticks (Budachetri et al., 2018; Duron et al., 2018; Rio et al., 2016; Adegoke et al., 2022; Gofton et al., 2015; Sassera et al., 2008). There are similarities in the bacteria identified from ticks in this study and those described from previous study. *Candidatus Midichloria, Rickettsia* and *Francisella* were amongst the dominant genera detected in ticks collected off migratory passerine birds from the neotropics moving into the United States (Budachetri et al., 2017). *Spiroplasma*, an endosymbiont well reported in dipteran vectors (Karatepe et al., 2018; Son et al., 2021) was also present in *Haemaphysalis* ticks in the current study. *Spiroplasma* was detected as a member of the *H. leporispalustris* microbiome, which could suggest an endosymbiotic relationship. While *Spiroplasma* species are well known for their male-killing attribute, several species of this symbiont do not induce male killing and present different phenotypes and genetic makeup, several of which benefit the host. We also detected *Candidatus Midichloria* in *I. brunneus*. *Candidatus Midichloria* is a ubiquitous endosymbiont of hard ticks first reported in *I. ricinus* (Sassera et al., 2006) and has been detected in several tick species.

Substantial differences in microbial diversity were not detected across tick genera, with *Amblyomma* having the lowest alpha diversity based on OTU abundance and the Shannon index. This low diversity is explained by the high relative abundance of *Francisella*. By applying PCoA analysis, we were able to visualize clustering in the microbial structure of tick genera, which illustrated several similarities and differences. Several tick species and genera share similar microbial profiles based on OTU abundance. Our findings also show that differences in the abundances of *Francisella* and most importantly *Rickettsia,* the two most abundant bacteria, were significant in shaping the PCoA cluster patterns. Our results show that ticks with high abundance of either *Francisella* or *Rickettsia* cluster separately from each other. BLAST analysis and phylogenetic reconstruction of these tick *Rickettsia* sequences demonstrated the presence of both spotted fever and non-spotted fever group *Rickettsia*. Several reports have detected *Rickettsia* in exotic ticks collected from birds migrating into North America, especially the USA (Miller et al., 2016; Souza et al., 2018; Budachetri et al., 2017) and Canada (Morshed et al., 2005; Ogden et al., 2015). However, the prevalence of these *Rickettsia* in ticks within the USA and the ability of these ticks to serve as competent vectors of *Rickettsia* remains poorly understood.

Our network analysis identified several unique microbial interactions across the different tick genera. First applied to the entire tick dataset, network analysis revealed more than 87% of interactions were positive, while more than half of the interactions were positive when applied to the individual datasets from each tick genus. We did, however, identify several environmental bacteria within networks consisting of tick endosymbionts and pathogens except in *Ixodes* ticks, for which we detected no pathogenic genus within the network. Of interest is the presence of *Rickettsia*, a pathogenic genus and several tick and non-tick symbionts within the same network, as seen in the *Haemaphysalis* dataset. An interesting observation was the significant positive correlation between *Rickettsia*, *Wolbachia*, and *Spiroplasma*, which suggests a potential interplay between these three bacteria genera. These observed differences in the microbial interactions detected across different tick gerea would suggest microbial interactions are uniquely shaped depending on the tick genus.

The findings from the current study support the hypothesis that migratory birds contribute to the seasonal introduction of non-native tick species and their associated microbes. Our analysis of the distribution of parasitized migratory birds suggests spring migration could facilitate dispersal hundreds to thousands of kilometers into the USA, with exotic ticks such as *Amblyomma coelebs, A. varium, A. parvum, A. nodosum, A. calcaratum* and several other ticks within this genus plausibly able to spread particularly far from where they were captured and sampled at stopover sites along the northern Gulf of Mexico.

## Conclusion

This study highlights the role migratory birds can play in introducing non-native tick species and their associated microbes into the USA. Our results demonstrate that songbirds have the ability to introduce exotic tick species during their seasonal migration into North America, with 421 ticks collected from 28 songbird species. Irrespective of the tick species, the core bacterial genera identified include *Francisella, Rickettsia, Spiroplasma,* and *Candidatus Midichloria*. The study provides valuable insight into the mechanisms of tick dispersal, emphasizing the importance of understanding bird migration patterns in predicting the introduction and establishment of potentially invasive tick populations. These findings contribute to the knowledge of the ecological factors influencing the spread of ticks and their associated pathogens, informing future strategies for surveillance and control efforts.

## Acknowledgments

We are especially thankful to the numerous University of Southern Mississippi field technicians who sampled birds. We appreciate the assistance of L. Calderón, W.C. Barrow, J.J. Buler, L.A. Randall, S. Collins, J. Smolinsky, A.N. Anderson, B. Geary, M. Dunning, and B.C. Wilson with various aspects of this study. We are grateful to the U.S. Fish and Wildlife Service, Louisiana Department of Wildlife and Fisheries, Alabama Department of Conservation and Natural Resources, The Nature Conservancy, The Louisiana Department of Culture, Recreation, and Tourism, as well as the Hollister, Guidry, and Nohr families for access to their properties and overall support. We thank J. Jenkins and B. Brown for feedback on an earlier version of this manuscript as well as anonymous reviewers for their critiques. This research was principally supported by a Pakistan-U.S. Science and Technology Cooperation Program award (U.S. Department of State), the Mississippi INBRE (an institutional Award (IDeA) from the National Institute of General Medical Sciences of the National Institutes of Health under award P20GM103476), and the NOAA RESTORE Act Science Program (NA17NOS4510092). The Alabama Ornithological Society provided additional funding. D. Becker was also supported by the National Science Foundation (BII 2213854). Any use of trade, firm, or product names is for descriptive purposes only and does not imply endorsement by the U.S. Government.

## Ethics Approval and Consent to Participate

The Institutional Biosafety Committee approved the protocol for the laboratory. All animal experiments were conducted per the guidelines in the Guide for Care and Use of Laboratory Animals of the National Institutes of Health, USA. The protocol for the collection and handling of birds was approved by the U.S. Geological Survey Bird Banding Laboratory (permit #24101), the Louisiana Department of Wildlife and Fisheries, the Alabama Department of Conservation and Natural Resources, and the Institutional Animal Care and Use Committee (IACUC) at the University of Southern Mississippi (protocol #17081101) and the U.S. Geological Survey (protocol #LFT 2019- 05).

## Consent of Publication

All authors read and approved the manuscript for publication.

## Availability of data and material

The datasets supporting the conclusion of this article are included within the article and its additional files. Raw data are available from the corresponding author upon request.

## Author contributions

Conceptualization: Shahid Karim, Lorenza Beati, Frank Moore

Data curation: Lorenza Beati, Deepak Kumar, Abdulsalam Adegoke, Shahid Karim, Theodore J. Zenzal Jr.

Formal analysis: Lorenza Beati, Deepak Kumar, Abdulsalam Adegoke, Daniel Becker, Shahid Karim

Funding acquisition: Shahid Karim, Frank Moore, Theodore J. Zenzal Jr.

Investigation: Lorenza Beati, Deepak Kumar, Theodore J. Zenzal Jr., Abdulsalam

Adegoke, Raima Sen, Latoyia Downs, Frank Moore, Shahid Karim

Methodology: Deepak Kumar, Raima Sen, Latoyia Downs, Lorenza Beati, Theodore J.

Zenzal Jr., Daniel Becker

Project administration: Shahid Karim, Theodore J. Zenzal Jr., Frank Moore

Resources: Shahid Karim, Theodore J. Zenzal Jr., Lorenza Beati, Frank Moore

Supervision; Shahid Karim, Theodore J. Zenzal Jr., and Frank Moore

Validation: Deepak Kumar, Shahid Karim

Visualization: Deepak Kumar, Abdulsalam Adegoke, Shahid Karim

Writing, original draft: Shahid Karim, Theodore J. Zenzal Jr., Abdulsalam Adegoke

Writing, review & editing: all authors

Any use of trade, firm, or product names is for descriptive purposes only and does not imply endorsement by the U.S. Government.

