## Supplementary material for "Ticks without borders: Microbial communities of immature Neotropical tick species parasitizing migratory landbirds along northern Gulf of Mexico": Fig. S1-S10

SUPPLEMENTARY FILES

List of reagents used in this study.

| Reagent name | Manufacturer | Location | Catalog Number |
| --- | --- | --- | --- |
| Dream Taq Polymerase | Invitrogen, Thermo Fisher Scientific | Grand Island, NY, USA | EP0701 |
| Sanger sequencing | Eurofins Genomics | Louisville, KY, USA |  |
| ZymoBIOMICS™ | Zymo Research | Irvine, CA, USA | D6306 |
| Bio-Rad | Bio-Rad | Hercules, CA, USA |  |
| Sigma-Aldrich | Sigma-Aldrich | St. Louis, MO, USA |  |

*Species-specific microbial interactions: Amblyomma*

Sixty-two percent of the total microbial interactions from the *Amblyomma* dataset were identified as positively correlated (Table S5). Due to the differences in the numbers from each *Amblyomma* species, they were divided into samples from a large sample size (> 20) and a small sample size (< 20). In the network analysis performed on the datasets with large sample sizes (*A. calcaratum, A. longirostre, A. maculatum, A. nodosum, A. parvum,* and *A. sabanerae*), we observed 56 positive and 46 negative significant partial correlations (Fig. S7, and Table S6). Several of the identified genera belonged to environmental groups and a few bacteria identified in other arthropod microbiomes such as *Wolbachia* and *Spiroplasma*. *Spiroplasma* was positively correlated with both *Wolbachia* and *Coxiella*. However, we did observe negative partial correlations between *Francisella* and *Rickettsia* similar to the whole tick dataset. *Candidatus Midichloria* was positively correlated with *Methylobacterium Methylorubrum* and *Escherichia Shigella*, which were both from environmental sources (Fig. S7). Of the datasets with small sample size (*A. americanum, A. coelebs, A. geayi, A. ovale, A. triste,* and *A. varium*), 25 microbial interactions were positive significant correlations, while 10 were negatively correlated (Fig. S8, and Table S7). From the network analysis, 5 unique clusters were identified, with the largest involving interactions between 10 bacteria genera. In contrast to the *Amblyomma* datasets with large representation, *Francisella* was not detected within any of the network clusters. In addition, *Coxiella* was positively correlated with *Sphingomonas,* while *Candidatus Midichloria* was negatively correlated with *Bacillus*.

*Species-specific microbial interactions: Ixodes*

Of the 54 significant partial interactions detected in the dataset from *Ixodes* ticks, 42 were positive interactions, while 12 were negative interactions (Tabel S8). From the network analysis, three different clusters were identified within the network analysis. However, we did not detect *Francisella* and *Rickettsia* in the *Ixodes* network dataset, indicating that other microbes, which include several of the bacteria identified as environmental group, are more important in driving the microbial interactions. We observed positive partial correlations between *Anaerobacillus, Wolbachia* and *Bacillus* and between *Wolbachia* and *Microbacterium* (Fig.S9, and Table S8). *Candidatus Midichloria* only interacted with *Lawsonella* and both show partial positive correlations.

*Species-specific microbial interactions: Haemaphysalis*

In the network analysis performed on *Haemaphysalis* ticks, we observed 48 significant partial positive correlations and 10 significant partial negative correlations which were grouped into three unique network clusters (Fig. S10, and Table S9). Two of the clusters involved commonly reported environmental group bacteria with previous associations to ticks and arthropods and were linked through direct or indirect interactions with *Rickettsia* (Fig. S10,). *Rickettsia* was positively correlated to *Spiroplasma* and *Candidatus Midichloria,* but negatively correlated with *Massilia*. *Candidatus Midichloria* was positively correlated to *Stenotrophomonas*, while *Stenotrophomonas* was also positively correlated to *Coxiella* (Fig. S10, and Table S9)*.*

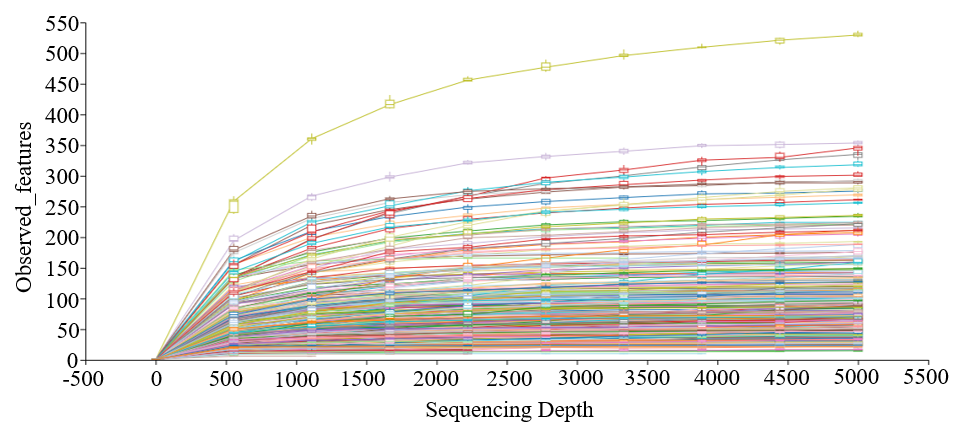

Figure S1: Rarefaction analysis of raw reads from all sampled ticks. Each curve represents an individual tick, and all tick sequences were normalized to a sequence depth of 5000.

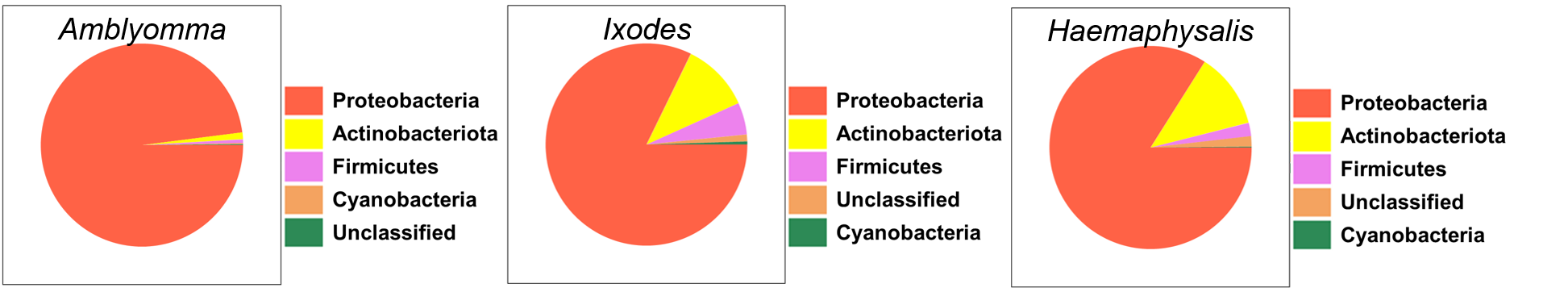

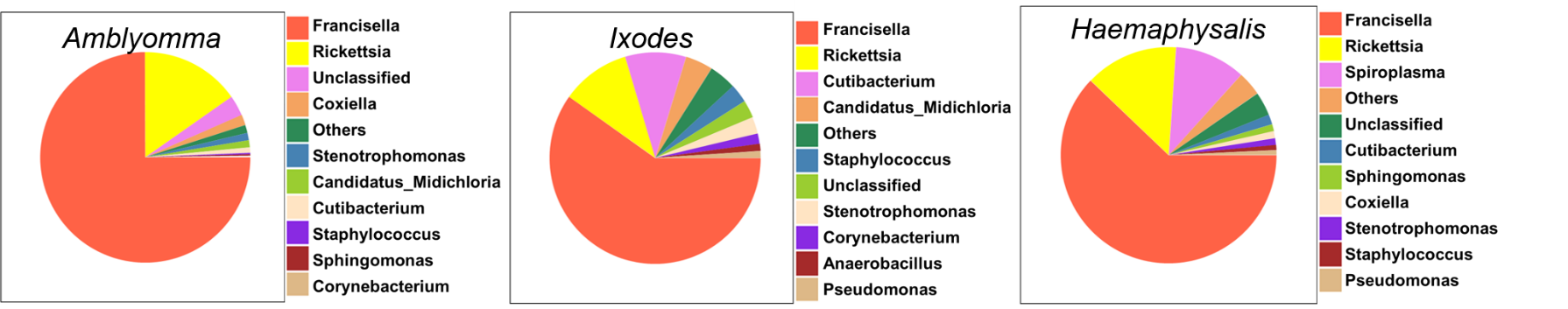

Figure S2: Pie chart summary of microbial abundances at A) phylum and B) genus level showing the taxa represented at an abundance of more than 1%.

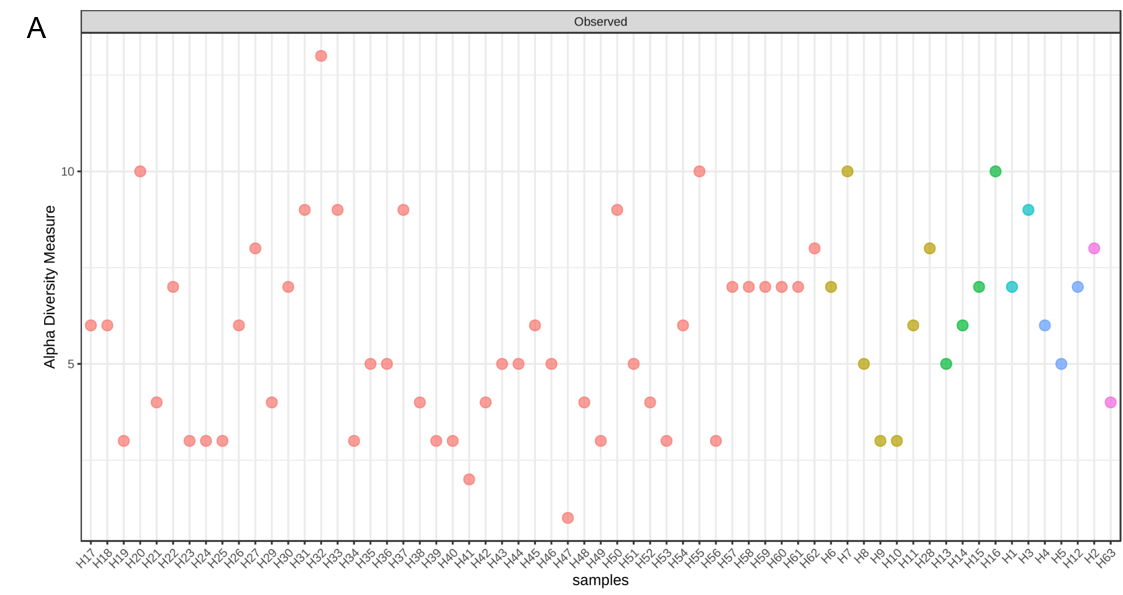

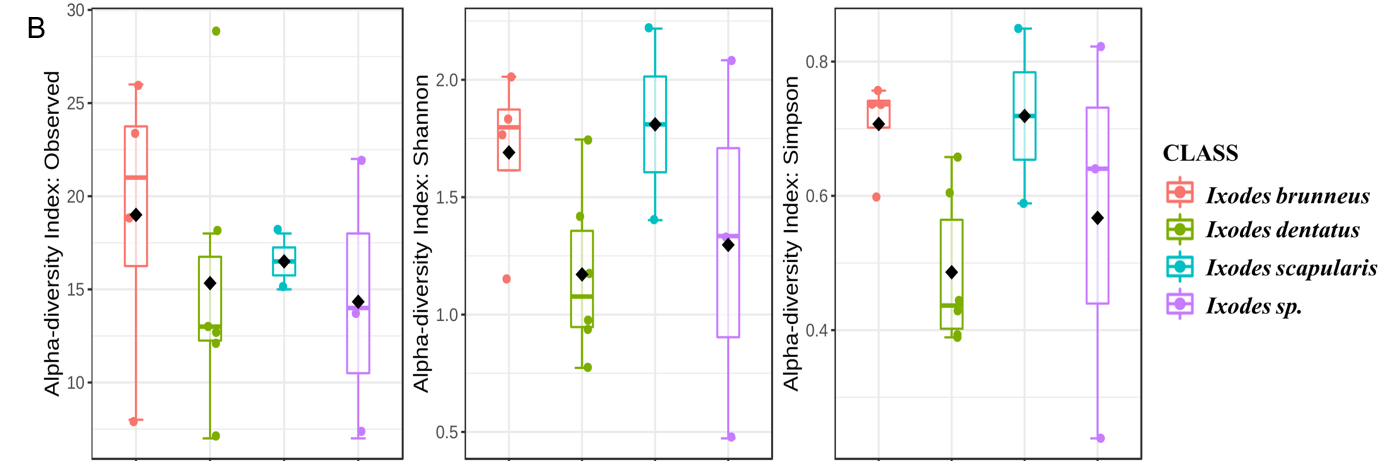

Figure S3: Alpha-diversity analysis of A) *Haemaphysalis leporispalustris* showing individual tick replicates and B) *Ixodes* ticks showing combined result. Alpha diversity was analyzed using the Observed OTUs, Shannon’s Index and Simpson’s Index metrics.

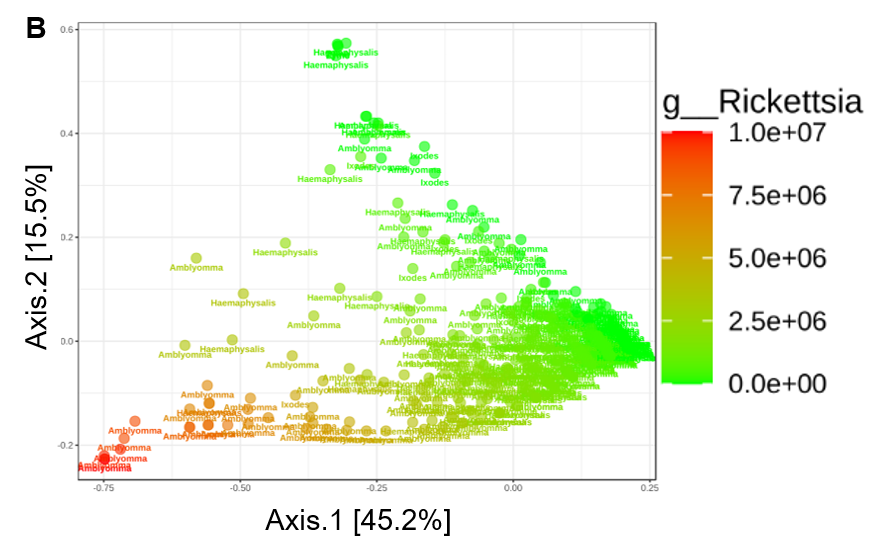

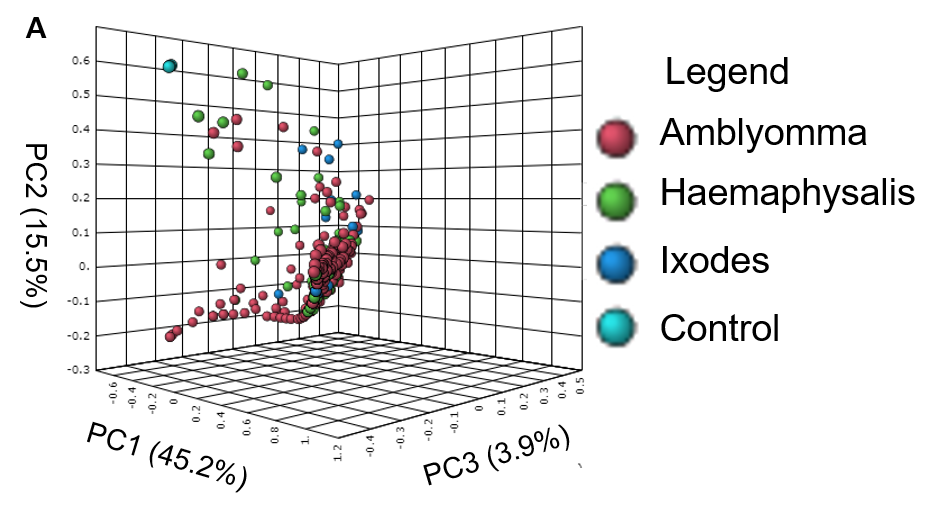

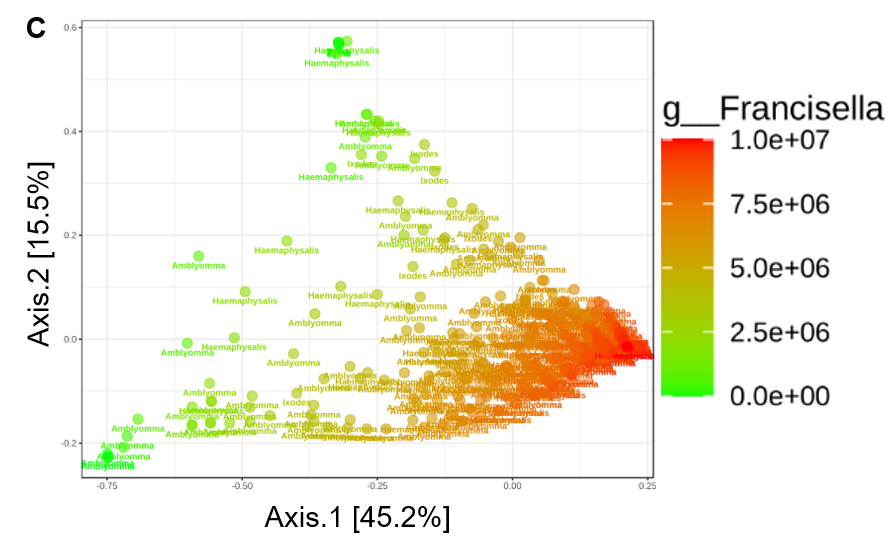

Figure S4: Principal coordinate analysis of beta-diversity measures for all tick dataset using the Bray-Curtis distance measures showing A) 3-D rendering of the PCoA. PCoA analysis based on taxon abundance showing the distribution of B) *Francisella* and C) *Rickettsia* and how the two genera shape the beta diversity.

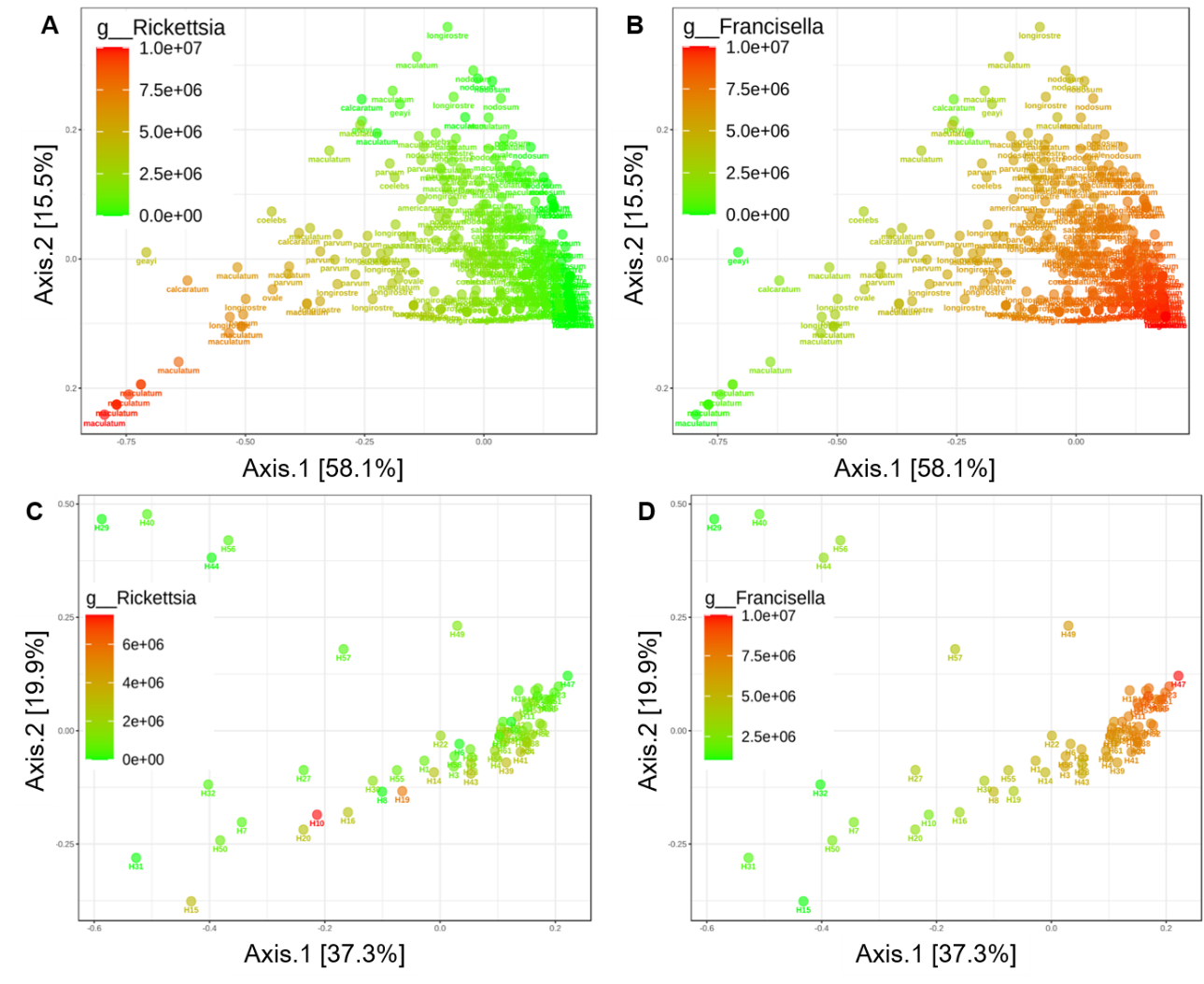

Figure S5: PCoA and taxon abundance distribution of *Rickettsia* and *Francisella* in A, B) *Amblyomma* and C, D) *Ixodes* ticks.

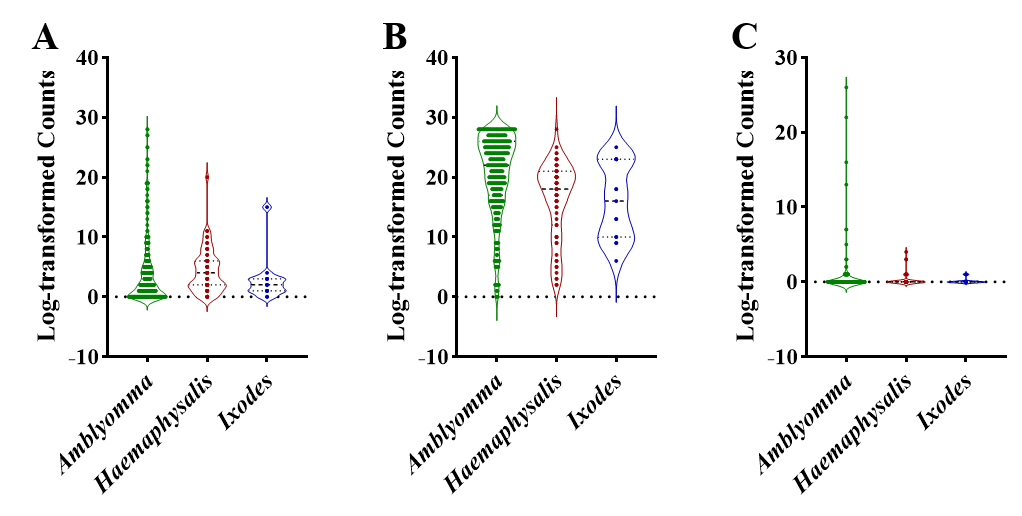

Figure S6: Violin plots of Log-transformed counts comparing the load of A) *Rickettsia,* B) *Francisella,* and C) *Coxiella* among *Amblyomma, Haemaphysalis,* and *Ixodes* ticks.

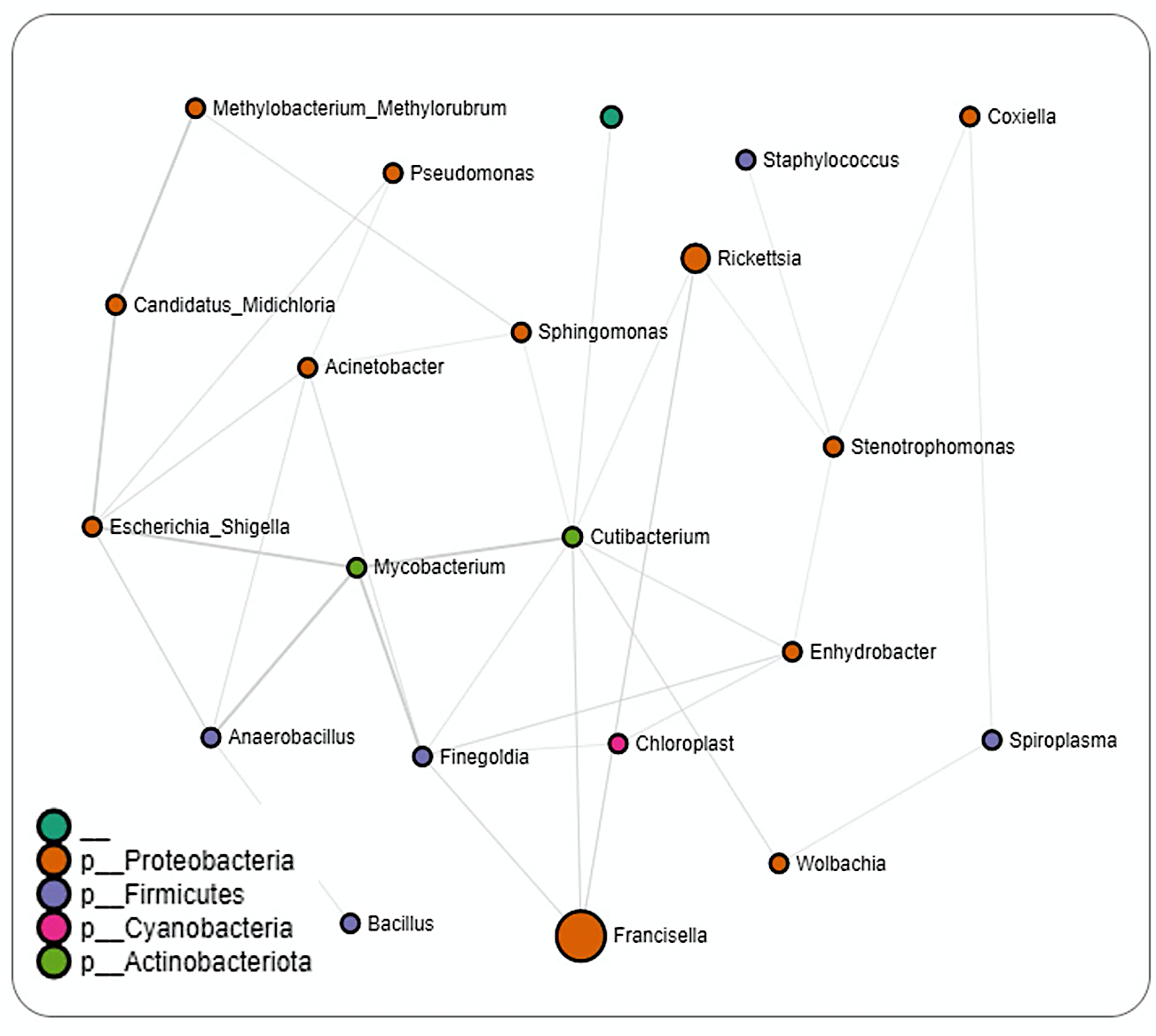

Figure S7: Correlation network analysis on *A. calcaratum, A. longirostre, A. maculatum, A. nodosum, A. parvum,* and *A. sabanerae*. Correlation network generated using the SparCC algorithm. Correlation network with nodes representing taxa at the family level and edges representing correlations between taxa pairs. Node size correlates with the number of interactions in which a taxon is involved. The color-coded legend shows the bacterial phyla. Nodes without a label represents unidentified taxa.

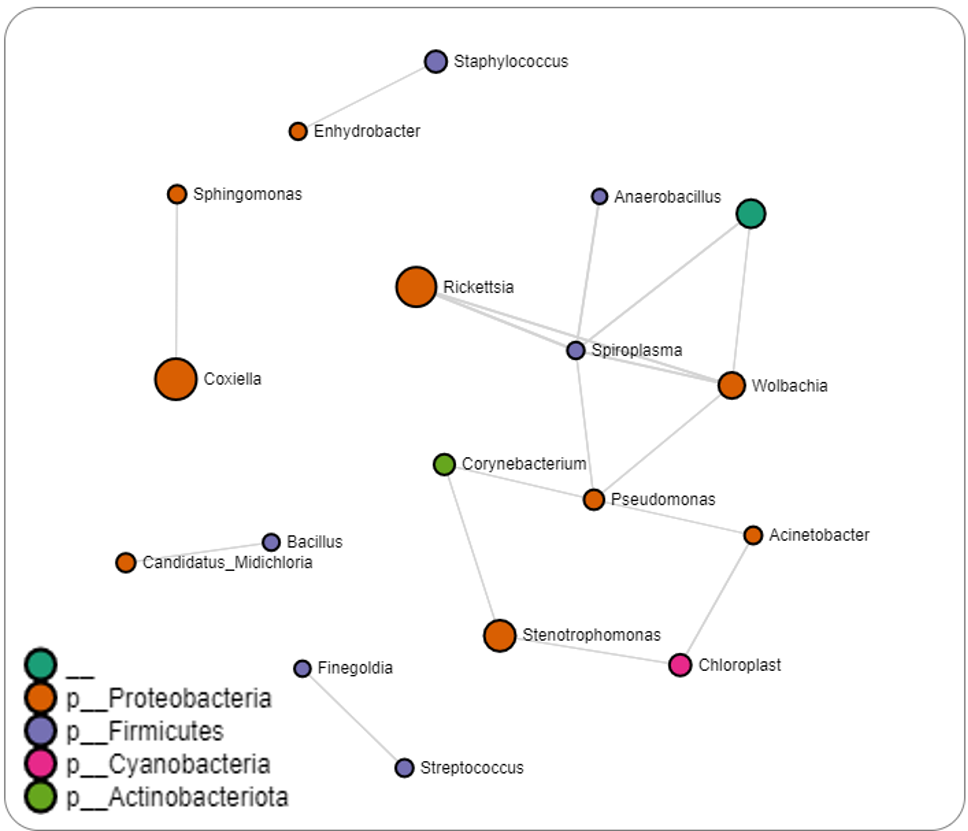

Figure S8: Correlation network analysis on *A. americanum, A. coelebs, A. geayi, A. ovale, A. triste,* and *A. varium*. Correlation network generated using the SparCC algorithm. Correlation network with nodes representing taxa at the family level and edges representing correlations between taxa pairs. Node size correlates with the number of interactions in which a taxon is involved. The color-coded legend shows the bacterial phyla.

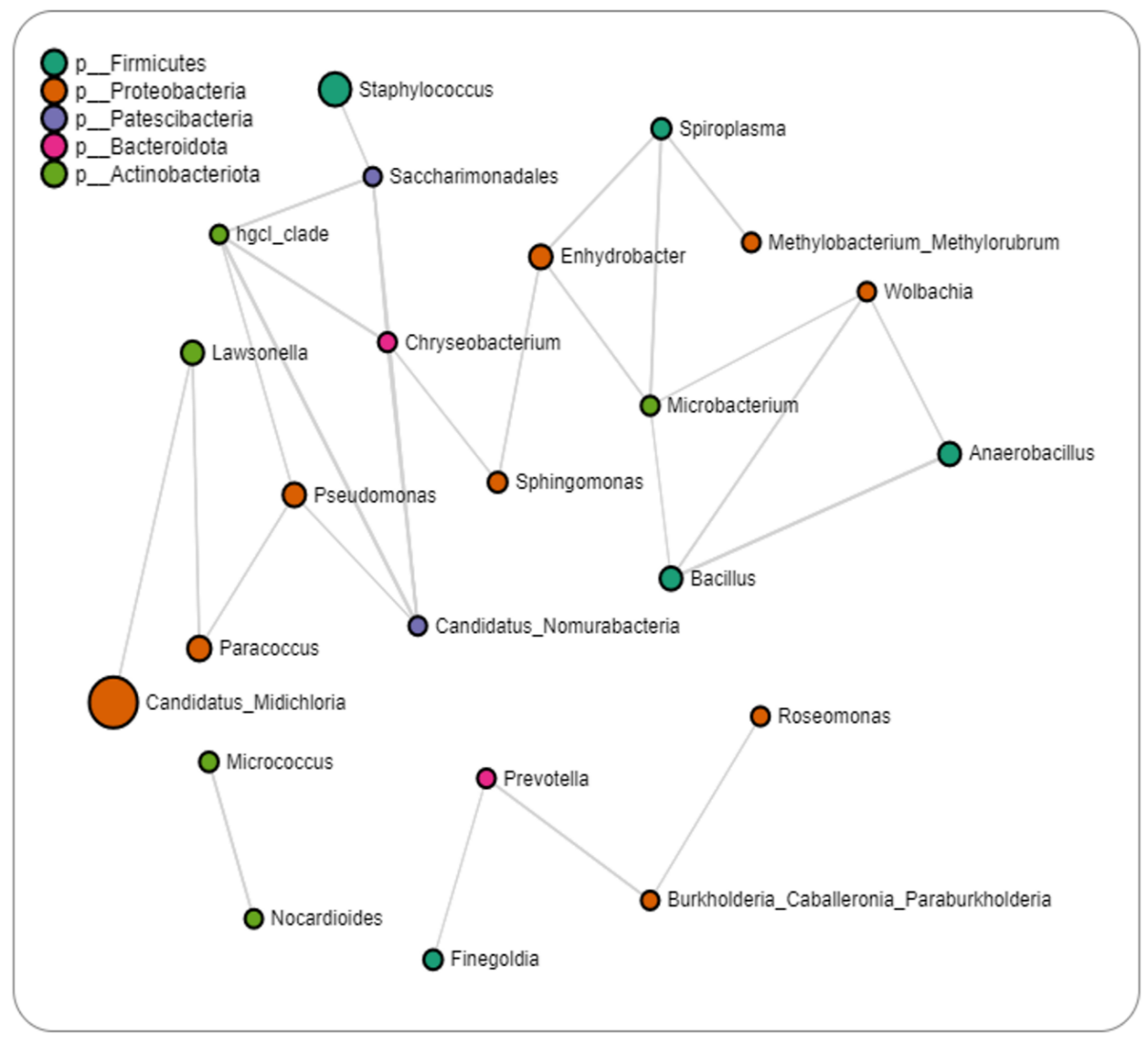

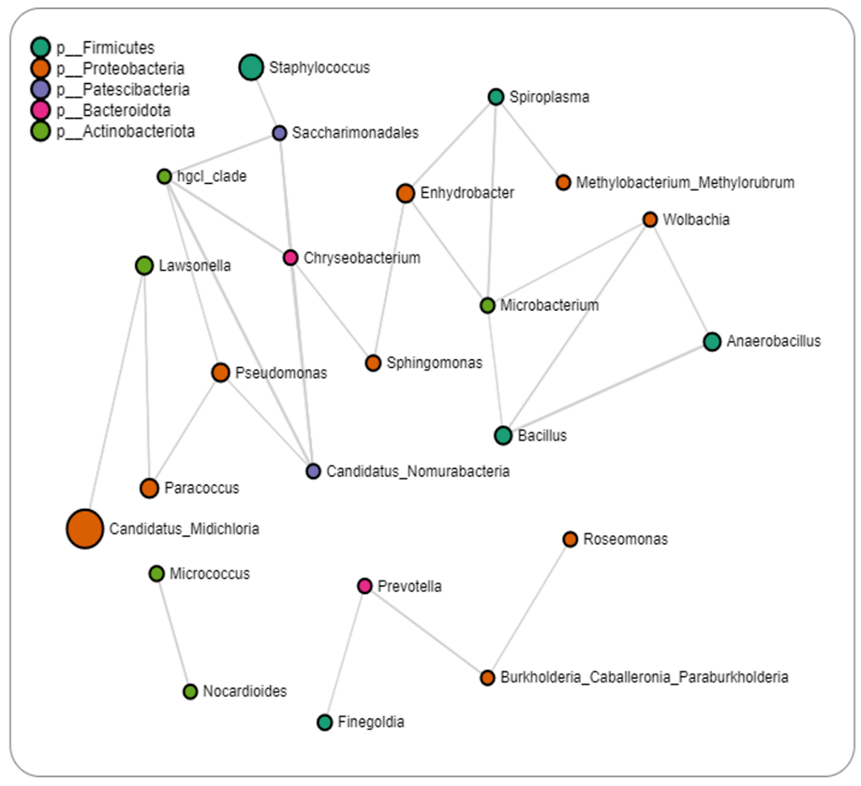

Figure S9: Correlation network analysis on *Ixodes* ticks. The correlation network generated using the SparCC algorithm. Correlation network with nodes representing taxa at the family level and edges representing correlations between taxa pairs. Node size correlates with the number of interactions in which a taxon is involved. The color-coded legend shows the bacterial phyla.

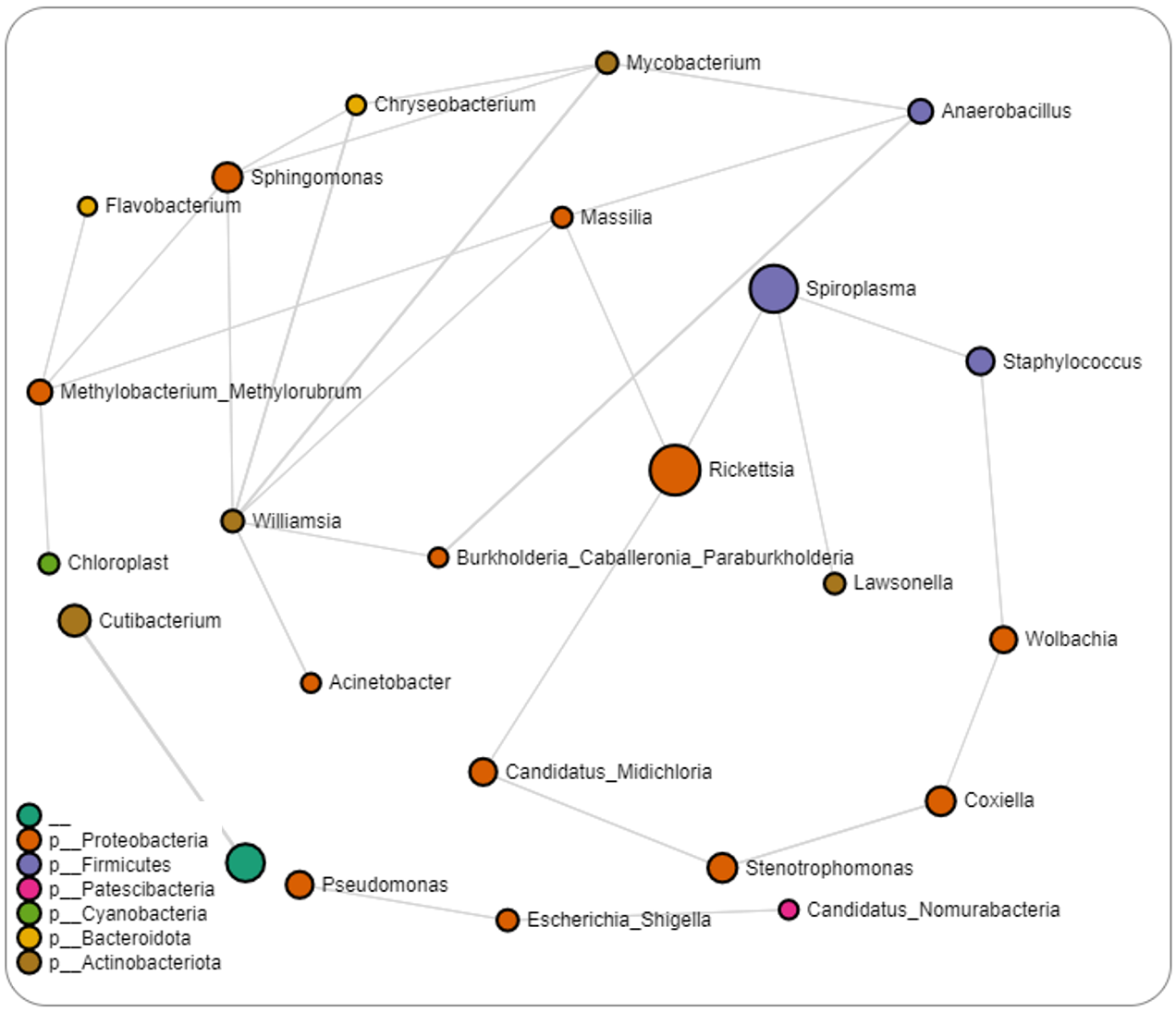

Figure S10: Correlation network analysis on *Haemaphysalis leporispalustris*. Correlation network generated using the SparCC algorithm. Correlation network with nodes representing taxa at the family level and edges representing correlations between taxa pairs. Node size correlates with the number of interactions in which a taxon is involved. The color-coded legend shows the bacterial phyla.
